# Differential disruptions in population coding along the dorsal-ventral axis of CA1 in the APP/PS1 mouse model of Aβ pathology

**DOI:** 10.1101/2022.07.15.500093

**Authors:** Udaysankar Chockanthan, Krishnan Padmanabhan

## Abstract

Alzheimer’s Disease (AD) is a progressive neurodegenerative disorder characterized by a range of behavioral alterations including memory loss as well as cognitive and psychiatric symptoms. While there is growing evidence that cellular and molecular pathologies, such as the accumulation of amyloid beta (Aβ) plaques may contribute to AD, it remains unclear how this histopathology can give rise to such disparate behavioral deficits. One hypothesis for these diverse behavioral presentations is that Aβ accumulation has differential effects on neuronal circuits across brain regions, depending on the diverse neurophysiological properties and connections within and between the neurons in these different areas. To test this, we recorded from large neuronal populations in the dorsal and ventral CA1 regions of the hippocampus, areas that are known to be structurally and functionally diverse, in both APP/PS1 animals, a mouse model of Aβ pathology, and age-matched C57BL/6 controls. Although we found similar levels of Aβ pathology in the two subregions, populations of neurons in dorsal and ventral CA1 in APP/PS1 mice showed distinct signatures of disrupted neuronal activity as animals navigated a virtual reality environment. In dorsal CA1, pairwise correlations and entropy, a measure of the diversity of activity patterns, were decreased in the APP/PS1 mice. However, in ventral CA1, the opposite findings were observed; pair-wise correlations and entropy was increased in APP/PS1 mice relative to C57BL/6 controls. When we attempted to connect the microscopic features of population activity (the correlations) with the macroscopic features of the population code (the entropy) using a pairwise Ising model, we found that the models’ performance decreased in predicting dorsal CA1 activity but increased in predicting vCA1 activity in APP/PS1 mice, as compared to the C57BL/6 animals. Taken together, these findings suggest that Aβ pathology exerts distinct effects across different hippocampal regions. Our results suggest that the diverse behavioral deficits associated with AD and the cellular pathology that arises from Aβ accumulation may be mechanistically linked by studying the dynamics of neural activity within these diverse hippocampal circuits.

## Introduction

Patients with Alzheimer disease (AD) experience an array of cognitive and behavioral symptoms, the most notable of which is memory loss (Greene et al., 1996; McKhann et al., 1984; Mega et al., 1996; Savva et al., 2009). Interestingly, a mix of additional neurologic and psychiatric symptoms, including deficits in spatial cognition (Delis et al., 1992; Henderson et al., 1989; Logsdon et al., 1998), agitation (Lyketsos et al., 2000; Steinberg et al., 2008), and elevated anxiety (Geda et al., 2008; Starkstein et al., 2007), suggest the disorder affects multiple neural circuits throughout the brain beyond those implicated solely in memory. Although the diagnosis of AD has conventionally been made by characterizing behavioral change in patients, considerable effort has been focused on identifying the molecular and cellular mechanisms that may underlie these deficits. This has, in part, been to identify potential targets to which treatment can be directed, but also to determine if an underlying pathology could account for the myriad symptoms experienced by patients.

The two major hallmarks of the disease, the accumulation of amyloid β (Aβ) (Braak and Braak, 1991; Hardy and Higgins, 1992; Thal et al., 2002) and tau neurofibrillary tangles (Braak and Braak, 1991; Braak and del Tredici, 2011; Mandelkow and Mandelkow, 1998) have been identified in both the neocortex and the hippocampus. Numerous studies have documented the impacts of this pathology on the structure and function of individual neurons. For example, Aβ pathology degrades dendritic spines (Tsai et al., 2004), and can lead neurons to be either hypoactive or hyperactive depending on their proximity to plaques (Busche et al., 2008). While studies at this level of analysis establish a critical link between histopathology and neuronal activity, they leave open the critical question of how microscopic changes at the cellular level lead to the constellation of symptoms in patients with AD.

Linking the pathology that occur due to Aβ or tau at the molecular and cellular level to the behavioral changes observed in patient and animal models remains a major challenge, especially in the context of identifying potential interventions (Cummings et al., 2014; Doody et al., 2014; Musiek and Bennett, 2021; Salloway et al., 2014; Sevigny et al., 2016). There are a number of reasons why this link remains an open area of inquiry. First, the behavioral heterogeneity seen in patients is not easily mapped to a unique cellular pathology. Second, a single pathology may affect the myriad neuronal circuits distributed throughout the neocortex and the hippocampus in different ways. Finally, if remains challenging to map on the diverse high-dimensional space that reflects the intrinsic cellular and synaptic properties of neurons (descriptors of the structure of the circuit) to the dynamics of population activity in the brain (a descriptor of the function of the circuit). Interestingly, as population neuronal activity has increasingly been shown to be a powerful predictor of behavior, it has proposed that an alternative model with which to investigate diseases is to investigate the changes in the dynamics of network activity that arise as a result of the aggregate effects of a pathology on the molecular, cellular and synaptic components of the network (Belmonte et al., 2004). In this framework, critical insights could be gained into the link between the molecular and cellular hallmarks of AD and the behavioral deficits seen in patients by examining the neural circuit and its functional dynamics.

A test of this hypothesis in an animal model requires two critical elements. First, the model brain region should be structurally and functionally diverse from the perspective of the circuit. Second, the brain region should exhibit significant Aβ pathology. We focused on the CA1 region of the hippocampus, as it satisfies these two conditions.

First the hippocampus serves a wide array of functions, including learning, memory, affective processing, and social cognition (Henke, 1990; Jimenez et al., 2018; Kim et al., 1976; Kjelstrup et al., 2002; Klüver and Bucy, 1937; Mackinnon and Squire, 1989; Moser et al., 1995; O’Keefe and Dostrovsky, 1971; Okuyama et al., 2016; Scoville and Milner, 1957; Zola-Morgan et al., 1992), all of which are typically affected in patients with AD. Interestingly, these diverse functions of the hippocampus appear to be physically separated along its longitudinal axis (Fanselow and Dong, 2010; Moser and Moser, 1998). While the dorsal hippocampus is implicated in episodic memory and spatial cognition (Moser et al., 1995), ventral hippocampus is believed to be involved in anxiety and social memory (Ciocchi et al., 2015; Jimenez et al., 2018; Okuyama et al., 2016). This differentiation across the dorsoventral hippocampal axis extends across multiple levels of organization, from gene expression(Mark S. Cembrowski et al., 2016; Thompson et al., 2008) to anatomical connectivity (Cenquizca and Swanson, 2007; Meira et al., 2018; Padmanabhan et al., 2019, 2016; Swanson and Cowan, 1977) to coding properties (Ciocchi et al., 2015; Jimenez et al., 2018; Kjelstrup et al., 2008; Okuyama et al., 2016). Multiple studies have shown, for example, that the scale of spatial representation coarsens from the dorsal to the ventral hippocampus (Jung et al., 1994; Keinath et al., 2014; Kjelstrup et al., 2008).

Second, the hippocampus is well recognized as a locus of damage in AD. The hippocampus has high levels of Aβ plaque accumulation as well as outright neuronal loss in patients with AD and mouse models of Aβ pathology (Braak et al., 2011; Ferreira et al., 2019; Gómez-Isla et al., 1996; Hsia et al., 1999; Whitesell et al., 2018). This damage extends to the functional and computational level, as was demonstrated by studies of hippocampal place cells, neurons that preferentially activate when an animal visits a particular region of its environment their place fields (O’Keefe, 1976; O’Keefe and Recce, 1993). Compared to age matched controls, dorsal hippocampal place cells in mouse models of Aβ pathology had larger place fields and were consequently less informative about an animal’s position (Cacucci et al., 2008; Cayzac et al., 2015). Such deficits likely contributed to the poor performance of the Aβ mouse models on tests of spatial memory (Cacucci et al., 2008; Cayzac et al., 2015). A recent study in the APP/PS1 model of Aβ pathology (Jankowsky et al., 2001) showed disruptions not only in the activity of individual neurons, but in the collective activity of the overall dorsal CA1 (dCA1) population (Chockanathan et al., 2020).

The hippocampus is thus an anatomically and functionally heterogeneous structure. However, most studies of hippocampal physiology, especially in the context of AD, have focused on the dorsal hippocampus (Cacucci et al., 2008; Cayzac et al., 2015; Chockanathan et al., 2020; Jun et al., 2020; Lin et al., 2022). Whether pathophysiological signature of AD in the ventral hippocampus is similar to that in dorsal hippocampus, or if the effects of AD pathology are highly contingent on the local properties of the brain region is largely unknown (Masurkar, 2018), and is critical to understanding how the elements of pathology at the molecular and cellular level influence behavior through the intermediary of the neural circuit. To address this question, and to test whether the impacts of Aβ pathology were divergent across the dorsoventral hippocampal axis, we performed recordings of large neuronal populations in dorsal and ventral CA1 (vCA1) in aged C57BL/6 and APP/PS1 mice. To quantify and compare the features of population activity across two strains, we calculated the pairwise correlation structure and the entropy and built maximum entropy models of the population. We found that Aβ pathology had a differential impact across dorsal and vCA1; while dCA1 collective activity was weakened in the APP/PS1 mice, the activity was enhanced in vCA1.

## Materials and methods

### Mice

All experiments were performed in accordance with regulatory standards and were approved by the Institutional Animal Care and Use Committee (IACUC) at the University of Rochester. Six male APP/PS1 mice (Strain #034832,The Jackson Laboratory, Bar Harbor, ME) and four male C57BL6/J mice (Strain #000664, The Jackson Laboratory) were included in the study. The APP/PS1 mice expressed a chimeric mouse/human amyloid precursor protein with the APP695swe mutation as well as a mutant human presenilin 1 PS1-dE9 (Jankowsky et al., 2001). At the time of recording, mice were 14 to 19 months of age. Surgeries and recordings were randomized to ensure that no systematic biases were introduced. Mice were housed in transparent cages on a 12h/12h light/dark cycle. All recording were performed in the light phase. Mice were not used for any previous experiments or procedures.

### Virtual reality setup

A one-dimensional virtual track was generated using the Virtual Reality MATLAB (VirMEn) toolbox based on previously published designs (Chockanathan and Padmanabhan, 2021; Gauthier and Tank, 2018; Meshulam et al., 2017). The virtual track was projected onto a curved board, which occupied 180° of the visual field of the mouse. A rotational encoder in the axel of the run-wheel transmitted wheel movement information to the computer, which updated the position of the animal in the virtual track accordingly. Upon reaching the end of the virtual track, mice received a small sweetened milk reward from a lick spout and their position was then updated to the beginning of the track. To minimize ambient light, sound and electromagnetic interference, the rig was enclosed in an electrically shielded box.

### Head fixing

Prior to surgery, animals were anesthetized using a 1-2% isoflurane mixture and placed in a stereotactic surgical rig. The scalp was resected, scored, and a craniotomy sites for dCA1 (coordinates relative to bregma: 2.5mm caudal, 1.5mm right) and vCA1 (coordinates relative to bregma: 3.05mm caudal, 3.15mm right) were marked. Next, the skull was scored and a metal ground pin and 3D-printed PLA headframe were attached to the skull using dental cement (Ortho-Jet Powder and Jet Liquid, Lang Dental Manufacturing Company, Wheeling, IL) and veterinary adhesive (Vetbond, The 3M Company, Maplewood, MN). Post-surgical analgesia was provided for 72h using a subcutaneous injection of 0.5-1.0 mg/kg slow-release buprenorphine, in accordance with IACUC protocols.

### Run training

Animals were given one day to recover following headframe implantation surgery. Subsequently, a 7-day training period was started to habituate the mice to the run-wheel, the virtual reality environment, and the lick spout (Warner and Padmanabhan, 2020). Each day, mice were head-fixed in the virtual environment and allowed to run on the wheel for one hour. No electrophysiological recordings were performed during this habituation phase.

### Craniotomy

Animals were anesthetized using a 1-2% isoflurane mixture. A craniotomy was performed over dCA1 and vCA1 using the sites marked during the headframe implantation surgery. A silicone sealant (Kwik-Cast, World Precision Instruments, Sarasota, FL) was applied over the craniotomy sites for protection. After the surgery, mice were allowed to recover for 12-18 hours in their home cages before electrophysiological recordings. This recovery period has previously been shown to not affect animal behavior (Warner and Padmanabhan, 2020).

### Electrophysiology

Extracellular voltage recordings were performed using an open source microfabricated silicon electrode array (Du et al., 2011; Yang et al., 2020) comprised of 128 recording channels spread across four shanks, each of which was spaced 50 µm apart. The probes were coated with a fluorescent dye (Alexa Fluor, Thermo Fisher Scientific, Waltham, MA) and connected to an Intan RHD 128-channel headstage and RHD USB interface board (Intan Technologies, Los Angeles, CA).

After the recovery period, animals were head-fixed into the run-wheel in the virtual reality rig. The electrode array was placed on a stereotactic frame and guided to the location of the craniotomies. For dCA1, the probe was lowered 1-1.2 mm from the brain surface and for vCA1, the probe was lowered 4-4.2 mm from the brain surface. Two days of recordings were performed for each mouse. For most animals, each day of recording comprised of one 1-2h session in dCA1 and one 1-2h session in vCA1. Signals were acquired at 30kHz in the 0.1-3500 Hz frequency band.

### Spike sorting

Preprocessing and spike sorting was performed using the open-source toolbox Kilosort 2 (Pachitariu et al., 2016). The widefield raw electrophysiology data were high passed at 500Hz, the median signal from all channels was subtracted from each stream, and correlated noise across all channels was removed. A set of template waveforms and spike times was generated and iteratively updated to reconstruct the original data set. The putative single units that resulted from this process were manually curated using the visualization software Phy 2 (Rossant et al., 2016). Based on the waveforms, amplitudes, and inter-spike interval distributions, units were preserved, eliminated, or merged. Units without clear refractory periods were eliminated. Additionally, the physical location of each unit in the brain was estimated based on the relative magnitude of its action potentials across different channels. After spike sorting, the data were imported into MATLAB 2019a (The MathWorks, Inc., Natick, MA) for analysis of population activity.

### Firing rate correlations and graph properties

For each unit, a time-dependent firing rate trace was obtained by calculating the number of spikes in a sliding 10ms window and converting the resulting time-series to a z-score. The correlation coefficient between each pair of traces was calculated and visualized as a correlation matrix. To ensure that the mean correlation values were not disproportionately weighted towards recording sessions with high neuron counts, a random subsample of 250 unit pairs was taken from each session. The matrices were converted to graph representations, where each neuron was represented by a node (Newman, 2003). A threshold was applied to the matrices, such that any correlation value that exceeded the threshold was preserved as an edge between the two corresponding nodes, while edges that fell below the threshold were discarded. The relative degree was calculated by summing the number of edges in the graph and dividing by the total number of possible edges between all the nodes in the graph. The clustering coefficient was based on node triplets, defined as three nodes that are connected to each other by either two edges (open triplet) or three edges (closed triplet). The clustering coefficient was defined as the proportion of all node triplets that are closed triplets. To ensure that the graphs being compared were of the same size, networks of 16-neuron subsamples were constructed for each recording session. Graph measures were calculated using the MATLAB Brain Connectivity Toolbox (Rubinov and Sporns, 2010).

### Entropy

Spiking activity for each unit was binned into 10ms non-overlapping bins, such that a value of 1 would be assigned to the bin if at least one spike occurred in that 10ms interval, and a value of 0 would be assigned if no spikes occurred. When this was done for all the units in a given animal, a matrix of binned activity was generated. A single column of this matrix thus represented the activity of all the neurons of the population within a 10ms window, which we refer to as a pattern (Chockanathan and Padmanabhan, 2021; de Ruyter van Steveninck et al., 1997; Schneidman et al., 2006). In a population of 10 units, there would thus be 2^10^ = 1024 possible binary patterns. The entropy of the population, which quantifies the diversity of observed patterns, can be described with the following formula, where *k* denotes a particular pattern and *p_k_* denotes the probability of that pattern:

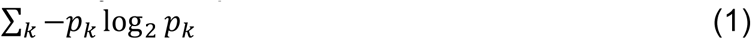

(de Ruyter van Steveninck et al., 1997). This number is then normalized by dividing by the mean firing rate of the population.

### Maximum entropy modeling

Fitting of maximum entropy models was performed using the maxent_toolbox (Maoz and Schneidman, 2017). The central principle of a maximum entropy model is the generation of a pattern probability distribution in which the activity of individual neurons and the coactivity of pairs of neurons match those of empirical probability distribution, but for which there is no further structure. More concretely, a maximum entropy model is a set of terms *h_i_*, which describe the average activity of individual neurons, and *J_ij_*, which describe the functional coupling between pairs of neurons. Given these terms, a pattern probability distribution for a neuronal population can be generated using the following:

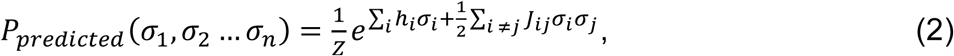

in which *σ_i_* denotes the binary state of neuron *i*, (*σ*_1_, *σ*_2_‖ *σ_n_*) denotes the predicted probability of a particular pattern, and Z indicates the partition function used to normalize the probability distribution. The objective is to iteratively adjust the terms *h_i_* and *J_ij_* such that they not only match the above constraints, but also result in a predicted pattern probability distribution with the least amount of structure (or alternatively, maximum entropy). The resultant predicted pattern probabilities thus represent the expected behavior of a population of neurons in the absence of any higher-order interactions beyond pairwise couplings (Meshulam et al., 2017; Schneidman et al., 2006).

To ensure sufficiently large populations, for the correlation, entropy, and maximum entropy calculations, only recording sessions with over 16 neurons were included. For each session, the neuronal population was repeatedly sampled to generate subpopulations of 16 neurons. The entropy, maximum entropy fit, and pattern probabilities of each subsample was calculated to generate a distribution of values for each recording session.

### Decoding brain region and strain identity from population activity

16-neuron subpopulations were generated for each recording session. For each subsample, the mean pairwise correlation, the entropy, and the KLD of the maximum entropy model were calculated; these were visualized in a 3D scatter plot. The data were then randomly split into training (80%) and testing (20%) subsets and used for three different classification tasks. The objective for the first task was to predict the brain region and strain identity (C57BL/6 dCA1 vs. APP/PS1 dCA1 vs. C57BL/6 vCA1 vs. APP/PS1 vCA1) for each point in the test set. For each point in the test set, the 5 nearest neighbors in the training set were identified; the mode identity of these 5 nearest neighbors was assigned as the predicted identity of the test set point. Accuracy was defined as the fraction of test points with correctly predicted identities. This process was repeated 100 times with different train/test set splits. A null distribution was generated by repeating this procedure after shuffling the identities of the points in the training set. The objective for the second and third tasks, respectively, was to predict the brain region for the C57BL/6 mice only (C57BL/6 dCA1 vs. C57BL/6 vCA1) and for the APP/PS1 mice only (APP/PS1 dCA1 vs. APP/PS1 vCA1).

### Tissue processing and staining

Transcardial perfusion was performed first with PBS solution, followed by a 4% paraformaldehyde (PFA) solution. The brain was removed from the skull, placed in a 4% PFA solution for 24 hours, then transferred to a 30% sucrose-PFA solution for 48 hours. Brains were frozen, sliced into 100µm coronal sections, mounted using Hoechst-containing media (Hoechst 33258, Invitrogen), and cover-slipped. Additionally, 26µm coronal sections spanning the anteroposterior extent of the hippocampus (approximately 1.25mm to 3.5mm posterior to bregma), spaced 400µm apart, were taken. These sections were stained for Aβ plaques with Congo Red (HT60-1KT, Milipore Sigma, Burlington, MA). Briefly, the sections were incubated in a 1:100 NaOH/NaCl solution for 20 minutes, stained with a 1:100 NaOH/Congo Red solution for 20 minutes, and rinsed several times with PBS before being mounted with Hoechst-containing media and cover-slipped.

### Slide imaging and scanning

The hippocampal sections stained with Congo Red were imaged using an Olympus VS120 slide scanner (Olympus Corporation, Tokyo, Japan).

### Aβ plaque quantification

The images were imported into an image analysis software Qupath (Bankhead et al., 2017), where Aβ plaque boundaries were manually circumscribed and hippocampal subfield boundaries were delineated. These data were then imported into MATLAB, where plaque numbers and fractional plaque area were computed for dorsal and ventral hippocampus and related to the electrophysiological properties of those respective regions.

## Results

To study the effects of Aβ pathology along the dorsal-ventral axis of CA1, we performed extracellular recordings in awake male C57BL/6 and APP/PS1 mice (age 14-19 months) in a virtual reality environment. Mice were head-fixed and placed on a non-motorized run-wheel. An image of a 1.9m virtual one-dimensional track was projected onto a curved screen in front of the mouse (Figure 1a, b), such that the movement of the wheel caused a corresponding change in the virtual track position (Chockanathan and Padmanabhan, 2021; Gauthier and Tank, 2018). When the mouse reached the end of the track, a sweetened milk reward was provided and the mouse was returned to the start of the track. After a 7-day period in which mice were habituated to the run-wheel and virtual environment, craniotomies were performed over dorsal and ventral CA1. Following a 12-18h recovery period, the animals were placed on the run-wheel and a 128- channel electrode array was lowered into dorsal or ventral CA1 (Figure 1c). Extracellular electrophysiological recordings lasting 1-2 hours were obtained over two days, allowing the identification of spikes (action potentials) from individual neurons (Figure 1d-g) while the animal was awake and behaving. There were no significant differences in running behavior between C57BL/6 and APP/PS1 mice as they navigated the virtual environment (Figure 1-figure supplement 1).

**Figure 1:**
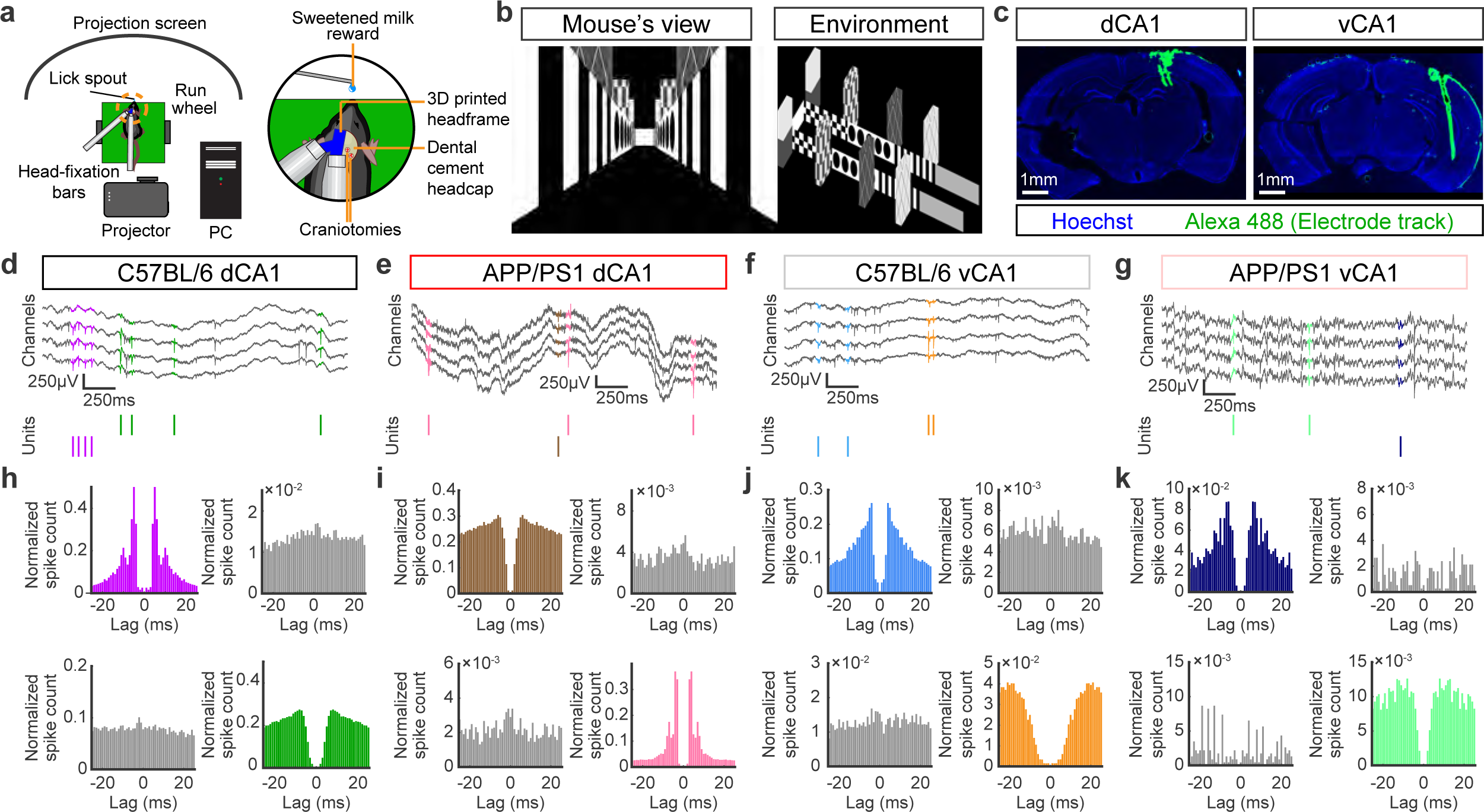
Experimental setup and spike sorting. (a) Left: schematic of head-fixed virtual reality recording rig. Right, close-up of mouse head, showing lick spout for sweetened milk reward delivery, headcap, and craniotomy sites. (b) Snapshots of virtual track, as seen from the mouse’s perspective and an overhead view. (c) Electrode arrays were coated with a fluorescent dye and targeted to either dorsal or ventral CA1. (d-g) Top: Example raw widefield electrophysiological traces in dorsal and ventral CA1 from C57BL/6 and APP/PS1 mice. In each trace, spikes from two different units are highlighted with unique colors. Bottom: Raster plot of spike times from the two units highlighted in the raw traces. (h-k) Auto-correlograms for each unit highlighted in panels d-g are shown with their respective colors. Cross-correlograms for each pair of units are shown in gray. Note the clear refractory periods in the auto-correlograms and the absence of such effects in the cross-correlogram.

From the band passed raw signals, we performed spike sorting across multiple neighboring channels using Kilosort 2 (Pachitariu et al., 2016) and the resulting putative units (neurons) were manually curated using Phy 2 (Rossant et al., 2016). Only units with clear refractory periods in their auto-correlograms (Figure 1h-k) and characteristic mean waveforms (Figure 2a-d) were included in subsequent analyses. Large populations of simultaneously recorded neurons were identified in dCA1 and vCA1 in both C57BL/6 and APP/PS1 mice (Figure 2e-h). From each dCA1 recording session, 33-133 units were identified in C57BL/6 mice and 26-44 units were identified in APP/PS1 mice. From the vCA1 recording sessions, an average of 61- 178 units were identified in C57BL/6 mice and 29-174 units were identified in APP/PS1 mice.

**Figure 2:**
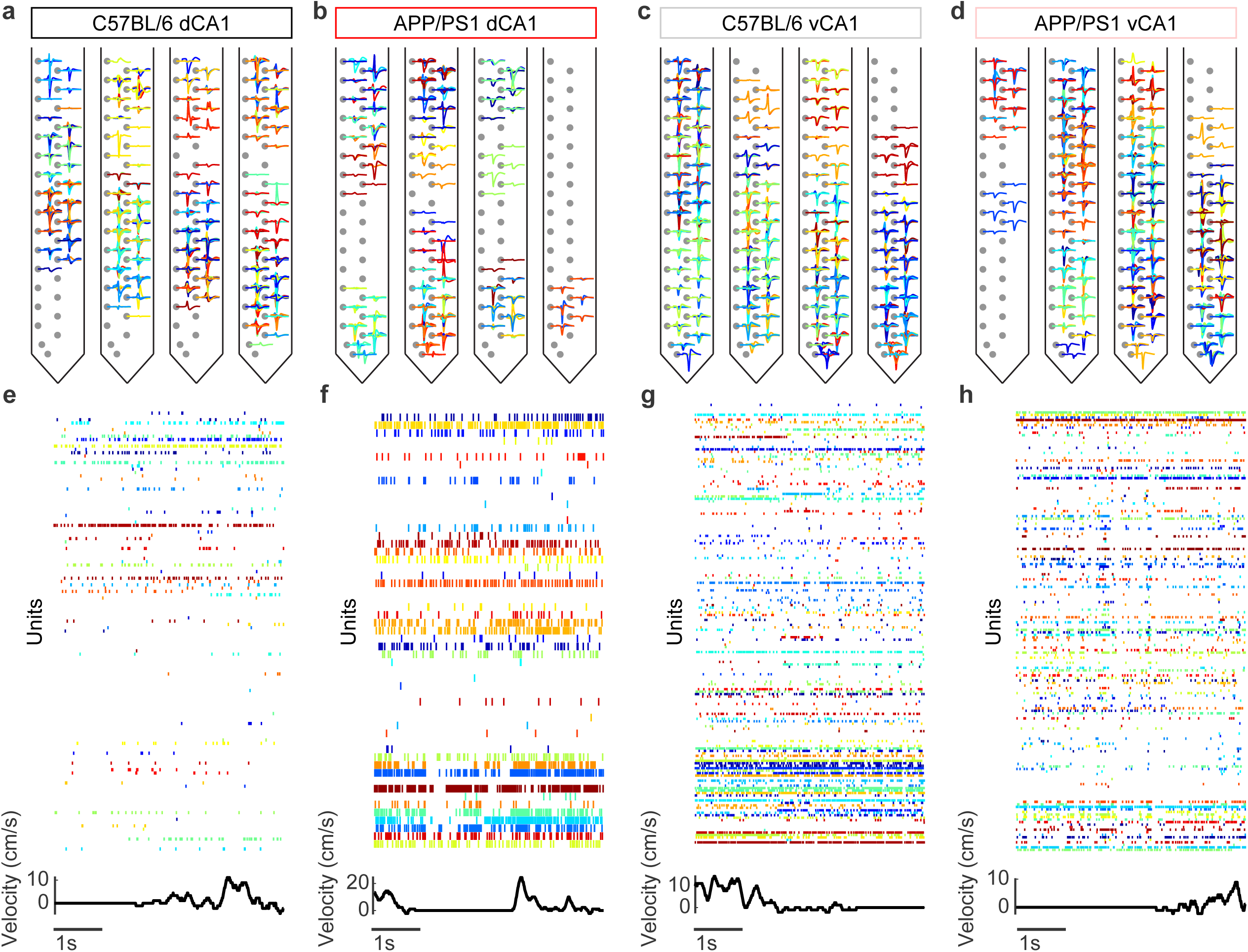
Identification of large populations of single units. (a-d) Mean waveform of all units in a representative recording session from dorsal and ventral CA1 in C57BL/6 and APP/PS1 animals. For each unit, waveforms are shown on the channel with the largest amplitude spike as well as up to four neighboring channels. (e-h) Top: Sample of population activity from each of the four groups, where the color of a given row on the raster plot refers to the mean waveform of that unit in panels a-d. Bottom: Simultaneous running velocity of the mouse.

After recordings were completed, mice were sacrificed and the brain was removed and sectioned to confirm the placement of the probe in either dCA1 or vCA1 and stained for Aβ plaques with Congo Red. Aβ plaques were identified throughout the dorsoventral axis of the hippocampus and cortex in APP/PS1 mice, but not in C57BL/6 animals (Figure 3a). Sections were digitized using a slide scanner and plaques were manually circumscribed. We found no significant difference in relative Aβ plaque area (Figure 3b) or density of Aβ plaques (Figure 3c) between dCA1 and vCA1. Furthermore, we found no association between the Aβ plaque burden and the mean firing rate of neurons in either dorsal or ventral CA1 (Figure 3d, e). This was consistent with earlier work demonstrating that the effects of Aβ pathology on neuronal activity could be heterogeneous in cortical neurons (Busche et al., 2008). It was therefore unlikely that we would be a measure such as a change in mean firing rate alone to be sufficient to capture the effects of Aβ on neural activity in APP/PS1 animals. Instead, previous studies have shown that a major and consistent feature of neural circuit disruption in models of Aβ pathology are the alterations in dendritic morphology (Šišková et al., 2014; Tsai et al., 2004), synapse density (Hsia et al., 1999; Tsai et al., 2004), and synaptic strength (Abramov et al., 2009a; Chapman et al., 1999; Palop et al., 2007). We thus reasoned that one consequence of Aβ pathology should be an alteration in the functional connectivity between neurons.

**Figure 3:**
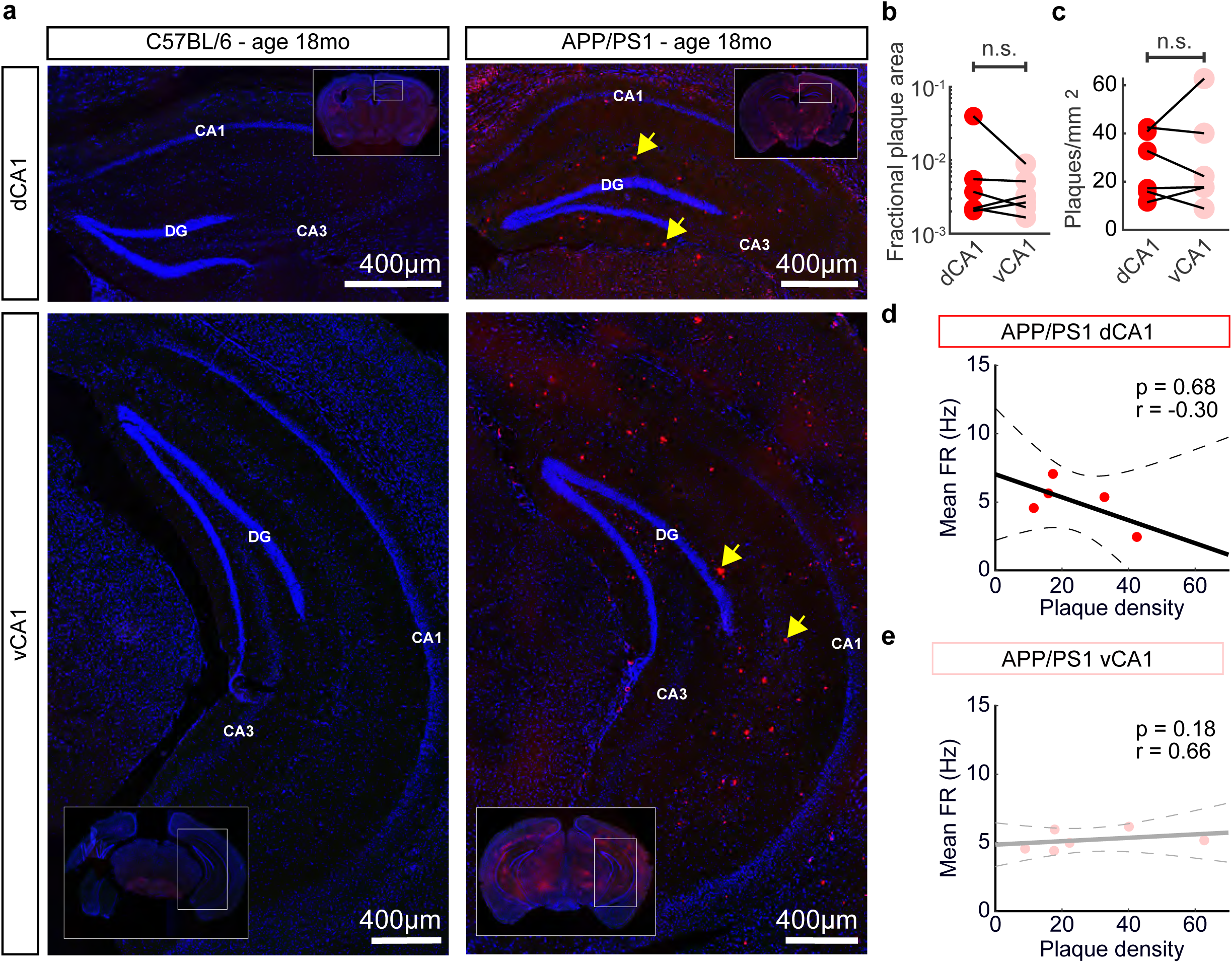
No significant differences in Aβ plaque burden between dCA1 and vCA1 in APP/PS1 mice. (a) Representative histological sections with from C57BL/6 and APP/PS1 mice with Congo Red staining for Aβ plaques (red) and Hoechst staining for cell nuclei (blue). Aβ plaques are present in both dorsal and ventral CA1 in APP/PS1 animals, but not in C57BL/6 animals. Yellow arrows indicate individual Aβ plaques. (b) There were no significant differences in fractional Aβ plaque area between dorsal and ventral CA1 in APP/PS1 mice (mean ± std: dCA1 = 9.2×10^-3^ ± 1.5×10^-2^, vCA1 = 4.0×10^-3^ ± 2.7×10^-3^, p > 0.99, two-sided Wilcoxon sign-rank test, n = 6 animals). Each point denotes a single animal. (c) There were no significant differences in plaque density between dorsal and ventral CA1 in APP/PS1 mice (mean ± std: dCA1 = 26.8 ± 13.6, vCA1 = 28.2 ± 19.8, p > 0.99, two-sided Wilcoxon sign-rank test, n = 6 animals). Each point denotes a single animal. (d) There was no significant correlation between the mean firing rate of dCA1 neurons and the dCA1 Aβ plaque density (r = -0.30, p = 0.68, Spearman rank correlation coefficient, n = 5 animals). Each point denotes a single animal. The black solid line denotes the least-squares regression and the black dashed lines denote the boundaries of the 95% confidence interval of the regression. (e) There was no significant correlation between the mean firing rate of vCA1 neurons and the vCA1 Aβ plaque density (r = 0.66, p = 0.18, Spearman rank correlation coefficient, n = 6 animals). Each point denotes a single animal. The grey solid line denotes the least squares regression and the grey dashed lines denote the boundaries of the 95% confidence interval of the regression.

To determine if such alterations in functional connectivity occurred across the CA1 region of hippocampus, we first calculated the correlations in activity between pairs of neurons (Figure 4a). Neuronal correlations have been implicated in a variety of cognitive processes, from spatial navigation (Meshulam et al., 2017; Skaggs et al., 1996; Stefanini et al., 2020) to attention (Mitchell et al., 2009; Rabinowitz et al., 2015), and disruptions in correlations have been associated with an array of neurologic disorders (Gonçalves et al., 2013; Hamm et al., 2017; Shuman et al., 2020). We found that pairwise correlations in dCA1 were decreased in APP/PS1 mice, relative to C57BL/6 mice (Figure 4b), consistent with earlier studies (Chockanathan et al., 2020). Interestingly, however, in vCA1, correlations in the APP/PS1 animals were larger than those in C57BL/6 animals (Figure 4c), suggesting a differential effect of Aβ pathology on the functional organization of neuronal connections across the dorsoventral hippocampal axis. These results were obtained using a temporal bin size of 10ms. However, the findings held true across a range of bin sizes up to 100ms (Figure 4-figure supplement 1).

**Figure 4:**
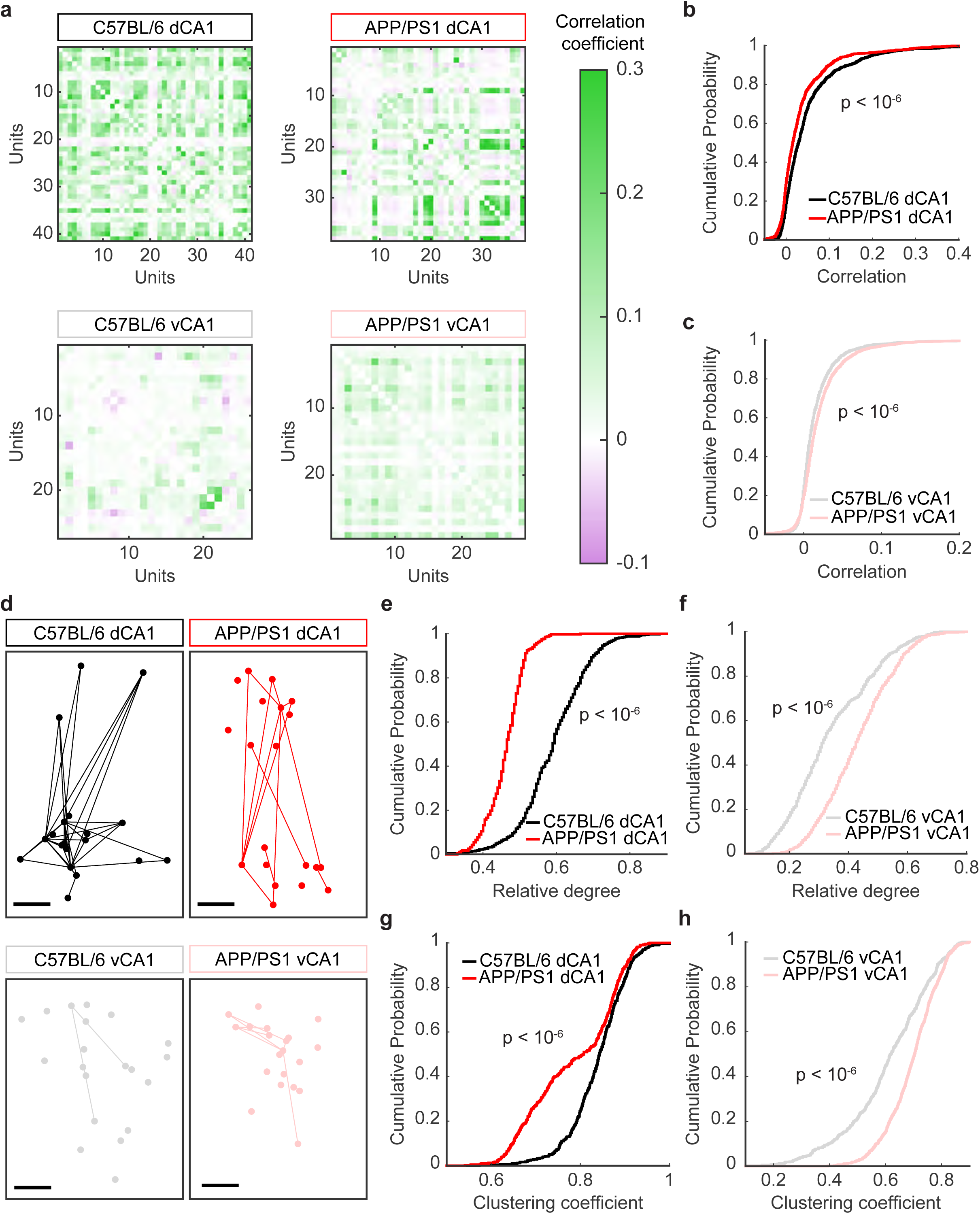
Pairwise correlations in APP/PS1 mice are decreased in dCA1, but increased in vCA1. (a) Matrices of correlation coefficients for all neurons in representative recording sessions. (b) In dCA1, mean correlations were lower in APP/PS1 mice than in C57BL/6 mice (mean ± std: C57BL/6 = 0.052 ± 0.075, APP/PS1 = 0.033 ± 0.063, p < 10^-6^, two-sided Wilcoxon rank-sum test, n_C57BL/6_ = 1250 unit pairs from 5 recording sessions, n_APP/PS1_ = 1000 unit pairs from 4 recording sessions). (c) In vCA1, mean correlations were higher in APP/PS1 mice than in C57BL/6 mice (mean ± std: C57BL/6 = 0.017 ± 0.031, APP/PS1 = 0.021 ± 0.034, p < 10^-6^, two-sided Wilcoxon rank-sum test, n_C57BL/6_ = 1500 unit pairs from 6 recording sessions, n_APP/PS1_ = 2000 unit pairs from 8 recording sessions). (d) Graph visualizations of neuronal populations. Circles denote neurons and lines denote correlations that exceeded a threshold of 0.06. The spatial position of each neuron corresponds to its approximate relative physical location. Scale bars denote 100µm. (e) In dCA1 graphs, the mean degree was smaller in APP/PS1 mice than in C57BL/6 mice (mean ± std: C57BL/6 = 0.57 ± 0.09, APP/PS1 = 0.45 ± 0.05, p < 10^-6^, two-sided Wilcoxon rank-sum test, n_C57BL/6_ = 500 samples from 5 recording sessions, n_APP/PS1_ = 400 samples from 4 recording sessions). (f) In vCA1 graphs, the mean degree was larger in APP/PS1 mice than in C57BL/6 mice (mean ± std: C57BL/6 = 0.32 ± 0.14, APP/PS1 = 0.40 ± 0.12, p < 10^-6^, two-sided Wilcoxon rank-sum test, n_C57BL/6_ = 600 samples from 6 recording sessions, n_APP/PS1_ = 800 samples from 8 recording sessions). (g) In dCA1 graphs, the clustering coefficient was smaller in APP/PS1 mice than in C57BL/6 mice (mean ± std: C57BL/6 = 0.83 ± 0.07, APP/PS1 = 0.77 ± 0.11, p < 10^-6^, two-sided Wilcoxon rank-sum test, n_C57BL/6_ = 500 samples from 5 recording sessions, n_APP/PS1_ = 400 samples from 4 recording sessions). (h) In vCA1 graphs, the clustering coefficient was larger in APP/PS1 mice than in C57BL/6 mice (mean ± std: C57BL/6 = 0.59 ± 0.16, APP/PS1 = 0.68 ± 0.10, p < 10^-6^, two-sided Wilcoxon rank-sum test, n_C57BL/6_ = 600 samples from 6 recording sessions, n_APP/PS1_ = 800 samples from 8 recording sessions).

As with metrics like firing rate, mean changes in pairwise correlations and functional connectivity represent the complex architecture of networks as a single value, which can often obscure critical aspects of network topology necessary to understand function. Inspired by studies from functional magnetic resonance imaging (fMRI) (Bullmore and Sporns, 2009; Chockanathan et al., 2019; Farahani et al., 2019) and previous work using graph theoretical approaches to study functional interactions across neuronal populations in CA1 (Chockanathan and Padmanabhan, 2021), we built on our correlation result by studying the topology of the functional connectivity we observed in our recordings. To do this, we constructed network graphs of the populations (Newman, 2003), in which nodes denoted neurons and edges denoted a pairwise correlation greater than 0.01 (Figure 4d). For each recording session, we took 500 16-unit subsamples of the overall population and calculated the average degree, or number of incident edges on each node, of the network. We found that in dCA1 networks, the average degree in APP/PS1 mice was smaller than that in C57BL/6 mice (Figure 4e). By contrast, in vCA1, the network degree in APP/PS1 mice was larger than that in C57BL/6 mice (Figure 4f). We found a similar result when calculating the clustering coefficient, which quantifies the extent to which the nodes of a network form densely connected groups. Dorsal CA1 networks in APP/PS1 animals were less clustered than those in C57BL/6 animals (Figure 4g), while vCA1 networks in APP/PS1 animals were more clustered than those in C57BL/6 animals (Figure 4h). Importantly, these differences in correlation and graph properties were not artifacts of mean firing rate variations, as there was no significant difference in firing rates between C57BL/6 and APP/PS1 mice in either dorsal or ventral CA (Figure 4 – figure supplement 2). Taken together, these findings demonstrate that the nature of network disruption in APP/PS1 mice was contingent on the brain region being studied, with dCA1 populations becoming more synchronous and vCA1 populations becoming less synchronous.

While pairwise correlations play a critical role in shaping the global patterns of activity (Meshulam et al., 2017; Schneidman et al., 2006), neuronal populations or ensembles exhibit complex collective behaviors that extend beyond just pairs of neurons (Ohiorhenuan et al., 2010). Previous studies have shown that studying ensemble activity can be critical for understanding sensory perception (Carrillo-Reid et al., 2019, 2016; Luczak et al., 2009; Miller et al., 2014), social cognition (Li et al., 2017; Liang et al., 2018), and spatial navigation (Meshulam et al., 2017; Stefanini et al., 2020). Additionally, two recent studies showed that changes in the statistical distributions of ensembles can serve as a marker for neuropsychiatric diseases (Chockanathan et al., 2020; Hamm et al., 2017). Ensemble patterns of activity across populations of neurons thus represent essential features of network function. We therefore sought to understand whether the properties of ensembles differed between C57BL/6 and APP/PS1 mice and how these disruptions varied across dorsal and ventral CA1.

One approach to do this is to simply quantify the distribution of patterns that are observed in the population. If we define a pattern as a combination of active neurons (represented as 1) and inactive neurons (represented as a 0) (Figure 5a), then in a 10-neuron population, there would be 2^10^ possible patterns. Each of these patterns can be thought of as a state of the network, and the probability distribution of the patterns thus reflects organization of the neuronal population. Some patterns will occur more frequently than others; for example, given the sparsity of neuronal activity, periods of silence, when all of the neurons are inactive, will occur more often than periods with large numbers of coactive neurons (Figure 5b). When we compared the probability distributions of patterns of activity between C57BL/6 and APP/PS1 mice, interesting themes emerged. In dCA1, the distributions were narrower, indicating a less diverse set of patterns in APP/PS1 mice as compared to C57BL/6 mice. By contrast, in vCA1, the pattern distributions were broader in APP/PS1 mice as compared to those seen in C57BL/6 mice (Figure 5b). One way to quantify these distributions is to calculate the entropy (see Materials and Methods). We found that patterns of activity in dCA1 were much more diverse in C57BL/6 animals, leading to a higher entropy as compared to the APP/PS1 animals (Figure 5c). Surprisingly, in vCA1, the entropy of the pattern distributions was higher in APP/PS1 mice as compared to C57Bl/6 mice, indicating a more diverse set of patterns (Figure 5d). As the entropy was normalized by the firing rate of the population, the resultant differences were not merely a consequence of differences in mean neuronal activity. Instead, these results reflected underlying differences in the number of different states or patterns that populations of neurons visited as the population activity evolved (Figure 5e,f). Finally, we found significant differences in entropy in dorsal and ventral CA1 between C57BL/6 and APP/PS1 animals across a range of subpopulation sizes (Figure 5-figure supplement 1). These data suggest that the diversity of patterns observed reflected an underlying principle of network organization across the hippocampus and that the organization is differentially affected in dCA1 versus vCA1.

**Figure 5:**
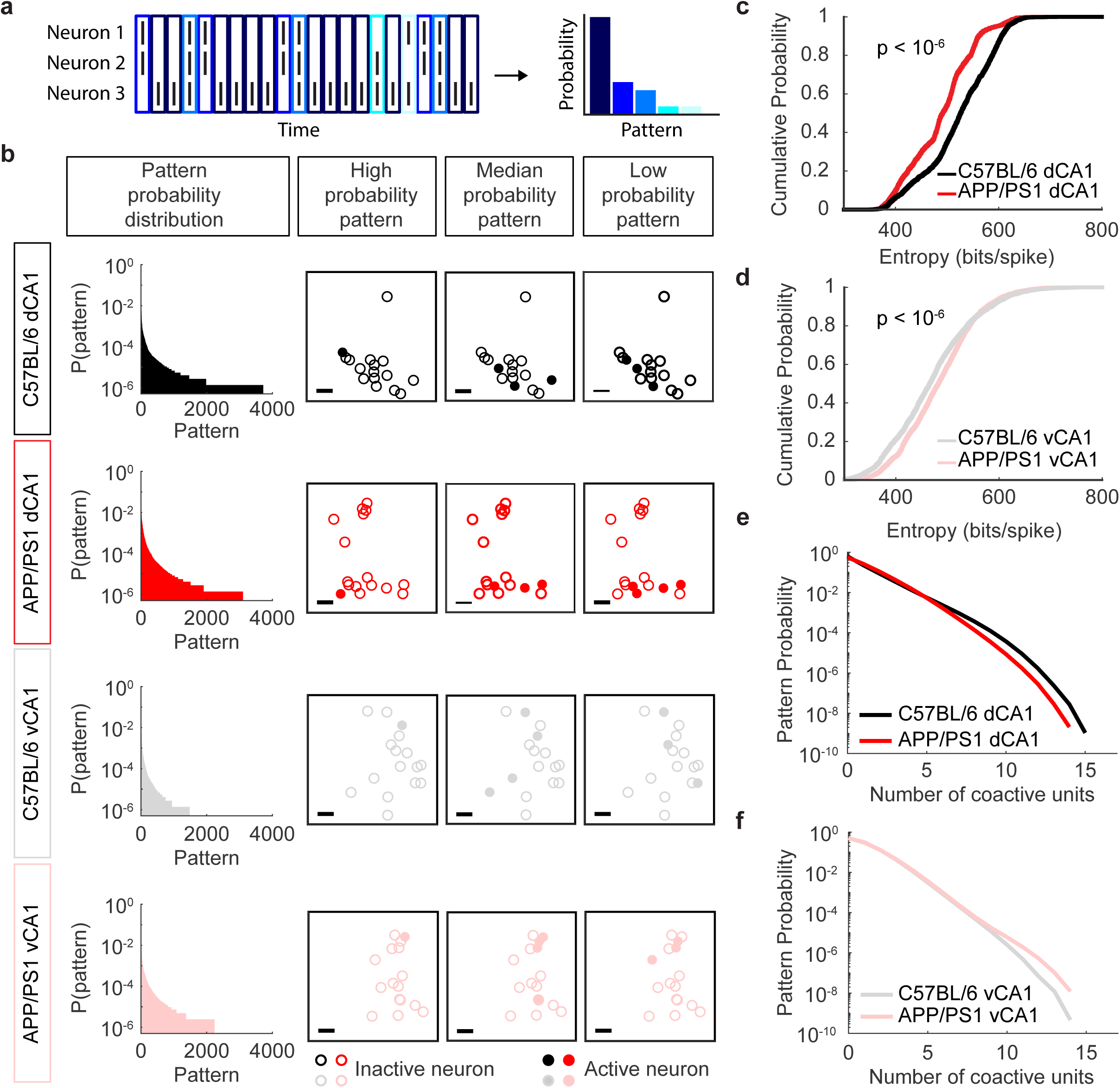
Population entropy in APP/PS1 mice is decreased in dCA1, but increased in vCA1. (a) Schematic of pattern identification and quantification using a 3-neuron cartoon population. A pattern is the combination of active and inactive neurons within a 10ms non-overlapping time window and can be conceptualized as a vertical column of the population spiking raster. In this 3-neuron population, there are 2^3^ possible patterns, though only 5 appear during this sample. Each pattern is assigned a unique shade of blue in the raster and in the probability distribution. (b) Left: representative examples of 16-neuron pattern probability distributions from dorsal and ventral CA1 in C57BL/6 and APP/PS1 mice. Right: visualizations of high, median, and low probability example patterns drawn from the corresponding probability distribution on the left. Open circles indicate inactive neurons and filled circles denote active neurons. The spatial position of each circle corresponds to its approximate relative physical position. Scale bars denote 100µm. (c) In dCA1 populations, the entropy was lower in APP/PS1 mice than in C57BL/6 mice (mean ± std: C57BL/6 = 522 ± 68 bits/spike, APP/PS1 = 489 ± 62 bits/spike, p < 10^-6^, two-sided Wilcoxon rank-sum test, n_C57BL/6_ = 2500 samples from 5 recording sessions, n_APP/PS1_ = 2000 samples from 4 recording sessions). (d) In vCA1 populations, the entropy was higher in APP/PS1 mice than in C57BL/6 mice (mean ± std: C57BL/6 = 471 ± 82 bits/spike, APP/PS1 = 484 ± 72 bits/spike, p < 10^-6^, two-sided Wilcoxon rank-sum test, n_C57BL/6_ = 3000 samples from 6 recording sessions, n_APP/PS1_ = 4000 samples from 8 recording sessions). (e) Probability distributions for dCA1 patterns grouped by number of coactive units. Pattern probabilities were averaged across 2500 subsamples from 5 recording sessions in C57BL/6 mice and across 2000 subsamples from 4 recording sessions in APP/PS1 mice. (f) Probability distributions for vCA1 patterns grouped by number of coactive units. Pattern probabilities were averaged across 3000 samples from 6 recording sessions in C57BL/6 mice and across 4000 samples from 8 recording sessions in APP/PS1 mice.

Calculations of entropy are an accounting of the macroscopic states or patterns generated by ensembles of neurons, while the time varying correlations shown in Figure 4 provide a microscopic detailing of the features of activity that might be constraining and shaping those states. We wanted to see if the pairwise interactions that reflect neuronal activity at the microscopic scale could be used to predict the macroscopic distributions of activity across the populations. To do this, we turned to a class of models called maximum entropy models, which aim to predict the distribution of patterns that occur across neural population while assuming as little as possible about the interactions governing that population (Meshulam et al., 2017; Ohiorhenuan et al., 2010; Schneidman et al., 2006; Shlens et al., 2006). In the second order models used here, the goal is to predict the probability distribution of patterns observed by considering only the activity (firing rate) of each neuron and the pairwise correlation between neurons. The model uses two sets of parameters: (1) a *h_i_* term for each neuron that describes its mean activity level and (2) a *J_ij_* term for every pair of neurons that describe their coactivity. Using these terms, we generated a synthetic probability distributions for all possible patterns and then compared the probabilities from the model to those measured in the data (Figure 6a). We visualized the synthetic and empirical probabilities using scatter plots, in which each point corresponded to a single pattern (Figure 6b), and we quantified the difference between the two probability distributions (data vs. model) using the Kulback-Leibler divergence (KLD). In dCA1, we found that the points in the scatter plots were closer to the unity line in APP/PS1 mice than in C57BL/6 mice (Figure 6b). This was reflected in the KLD, which was smaller in APP/PS1 mice, indicating a better agreement between the empirical and predicted pattern probabilities, than in C57BL/6 mice (Figure 6c). In vCA1, however, the KLD was higher, indicating a worse fit in the APP/PS1 animals than in the C57BL/6 animals (Figure 6d). These results, held true over a range of subpopulation sizes (Figure 6-figure supplement 1) and showed that, in APP/PS1 mice, the functional interactions that shape macroscopic patterns of neuronal activity were differentially affected in dorsal and ventral CA1. Consistent with previous findings (Chockanathan et al., 2020), relative to C57BL/6 mice, pairwise interactions were better able to predict the activity of the overall population in APP/PS1 mice in dCA1. On the other hand, activity patterns in APP/PS1 populations in vCA1 were less determined by those same pairwise interactions. Taken together, these results suggest that not only are the macroscopic and microscopic elements of network activity across the CA1 region of hippocampus differentially affected in the APP/PS1 model, but that the relationship between the microscopic interactions that govern correlations and the macroscopic features of global activity and network organization are altered by Aβ pathology.

**Figure 6:**
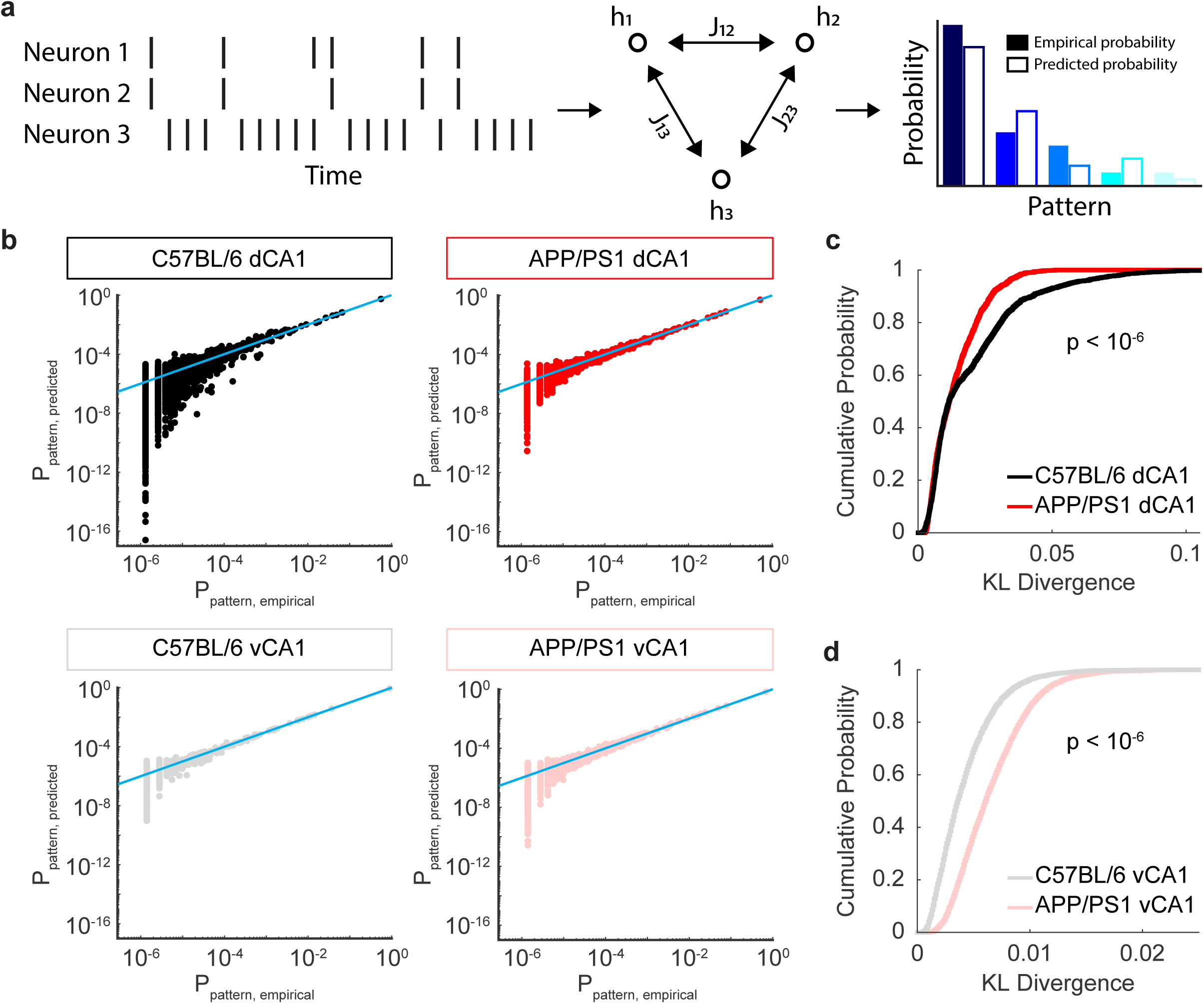
Prediction of pattern probabilities by pairwise interactions in APP/PS1 mice is improved in dCA1, but worsened in vCA1. (a) Schematic of maximum entropy model. From the population spike rasters, two sets of terms were fitted: a set of activity terms *h_i_* and a set of pairwise interaction terms *J_ij_*. From these terms, a synthetic pattern probability distribution was generated in which the mean activity of each neuron and the coactivity of every pair of neurons was identical to that of the empirical distribution but was otherwise unstructured. The predicted and empirical pattern probabilities were then compared. (b) Representative examples of maximum entropy model predicted probabilities and empirical probabilities. Each point denotes a single pattern and the grey line denotes unity. (c) In dCA1 populations, the KLD was lower in APP/PS1 mice than in C57BL/6 mice (mean ± std: C57BL/6 = 1.9×10^-2^ ± 1.8×10^-2^, APP/PS1 = 1.4×10^-2^ ± 9.3×10^-3^ bits/spike, p < 10^-6^, two-sided Wilcoxon rank-sum test, n_C57BL/6_ = 2500 samples from 5 recording sessions, n_APP/PS1_ = 2000 samples from 4 recording sessions). (d) In vCA1 populations, the entropy was higher in APP/PS1 mice than in C57BL/6 mice (mean ± std: C57BL/6 = 4.3×10^-3^ ± 2.7×10^-3^, APP/PS1 = 6.6×10^-3^ ± 3.3×10^-3^, p < 10^-4^, two-sided Wilcoxon rank-sum test, n_C57BL/6_ = 3000 samples from 6 recording sessions, n_APP/PS1_ = 4000 samples from 8 recording sessions).

The KLD between the predicted and empirical pattern distributions provided a measure of the error of the maximum entropy model. However, it is averaged over a diverse array of population patterns. Different patterns had different numbers of co-activated neurons (Figure 5b). To understand whether certain categories of patterns were predicted less accurately by the maximum entropy models than others, we examined patterns by the number of coactive neurons in both CA1 subfields in C57BL/6 and APP/PS1 mice. First, we found that the gap between the empirical and predicted probabilities for patterns with many coactive neurons was higher than those with few active neurons (Figure 7a). To dissect how these variations in prediction error related to the KLD, we calculated the total prediction error for each pattern category, where category was defined by the total number of coactive neurons, regardless of the specific combinatorial pattern of that activity. Across both strains of mice and in both CA1 subfields, the total prediction error was greatest for patterns with 4-5 coactive neurons. This was because the prediction error for patterns with 4-5 was relatively poor *and* there were more combinatorial patterns with 4-5 coactive neurons than there were for 15-16 coactive neurons. Thus, even though the prediction error of individual patterns increased with the number of coactive neurons (Figure 7-figure supplement 1), the largest contribution to the KLD came from the prediction error of patterns with 4-5 coactive neurons. For these patterns, and for most of the other categories, the total prediction error was lower in the APP/PS1 animals than the APP/PS1 animals in dCA1 (Figure 7b). This result was consistent with the decreased KLD in APP/PS1 mice in dCA1 (Figure 6c). In vCA1, the total prediction error curve was increased in APP/PS1 mice, relative to C57BL/6 mice (Figure 7c), consistent with the elevated KLD in APP/PS1 mice in vCA1 (Figure 6d). These findings indicate that in both dorsal and ventral CA1, the disruptions to population activity in APP/PS1 mice primarily arose from alterations in the coactivity of ensembles of 4-6 neurons, suggesting that small networks of cells were most sensitive to the effects of Aβ pathology. While the pattern prediction error was decreased in dCA1 in the APP/PS1 animals, it was increased in vCA1, suggesting that the effects of Aβ pathology percolated across the hippocampus in different ways.

**Figure 7:**
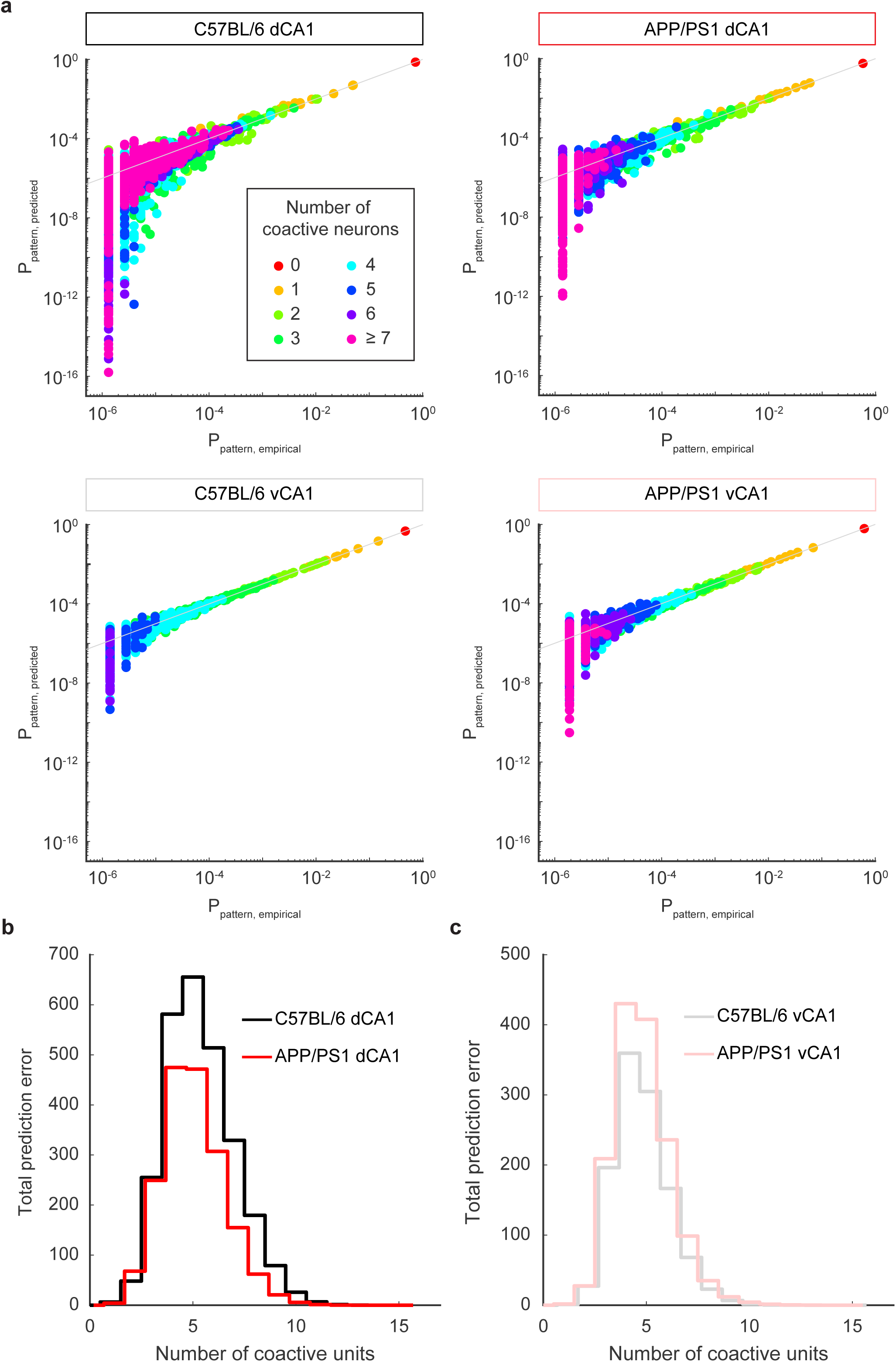
Disruptions in APP/PS1 population activity arise from disruptions to patterns with large numbers of coactive neurons. (a) Representative examples of maximum entropy model predicted probabilities and empirical probabilities. Each point denotes a single pattern and the grey line denotes unity. The color of the point corresponds to the number of coactive neurons in the pattern. (b) In dCA1, the total prediction error for APP/PS1 mice was lower than that for C57BL/6 mice across almost all pattern categories. The prediction error for patterns with 4-5 coactive units contributed most to the overall prediction error. Pattern probabilities were averaged across 2500 subsamples from 5 recording sessions in C57BL/6 mice and across 2000 subsamples from 4 recording sessions in APP/PS1 mice. (c) In vCA1, the total prediction error for APP/PS1 mice was higher than that for C57BL/6 mice across almost all pattern categories. The prediction error for patterns with 4-5 coactive units contributed most to the overall prediction error. Pattern probabilities were averaged across 3000 subsamples from 6 recording sessions in C57BL/6 mice and across 4000 subsamples from 8 recording sessions in APP/PS1 mice.

We have thus far examined the population code from three perspectives. The pair-wise correlations capture the interactions between the individual neurons, the entropy represents the diversity of patterns across the population, and the maximum entropy models reveal the extent to which the features of the network can be used to predict the global structure of the population. The distributions of each of these three properties varied between dCA1 and vCA1 as well as between C57BL/6 and APP/PS1 mice (Figure 8a). One can think of each of these measures as a feature of a high dimensional description of the population code. Each can then be represented as a single dimension in a coding space. Thus, plotting the mean correlation, the entropy, and the KLD of the maximum entropy model for each ensemble of neuronal activity for an ensemble population of 16 neurons (black: C57BL/6 dCA1, gray: C57BL/6 vCA1, red: APP/PS1 dCA1, pink: APP/PS1 vCA1) allows us to see the relative position of the neural population code within this space (Figure 8b). This revealed key insights. First, in the C57BL/6 mice, the dCA1 and vCA1 population clusters occupy different portions of the space, suggesting that the structure of the population code in dorsal and ventral CA1 is fundamentally different. Our analyses therefore reveal the consequences of the diverse anatomical and physiological properties of networks and circuits on the population code along the dorsal ventral axis of CA1 (Fanselow and Dong, 2010). Perhaps more striking, however, was the effect that Aβ pathology has on this population code. The networks in the APP/PS1 animals resided in an entirely new part of the coding space, suggesting that Aβ pathology fundamentally altered all aspects of the population code. Second, the effects of Aβ pathology on population activity depended on whether that population was in dCA1 or vCA1. Finally, the dorsal and ventral CA1 clusters were closer to one another in the APP/PS1 mice than in the C57BL/6 animals.

**Figure 8:**
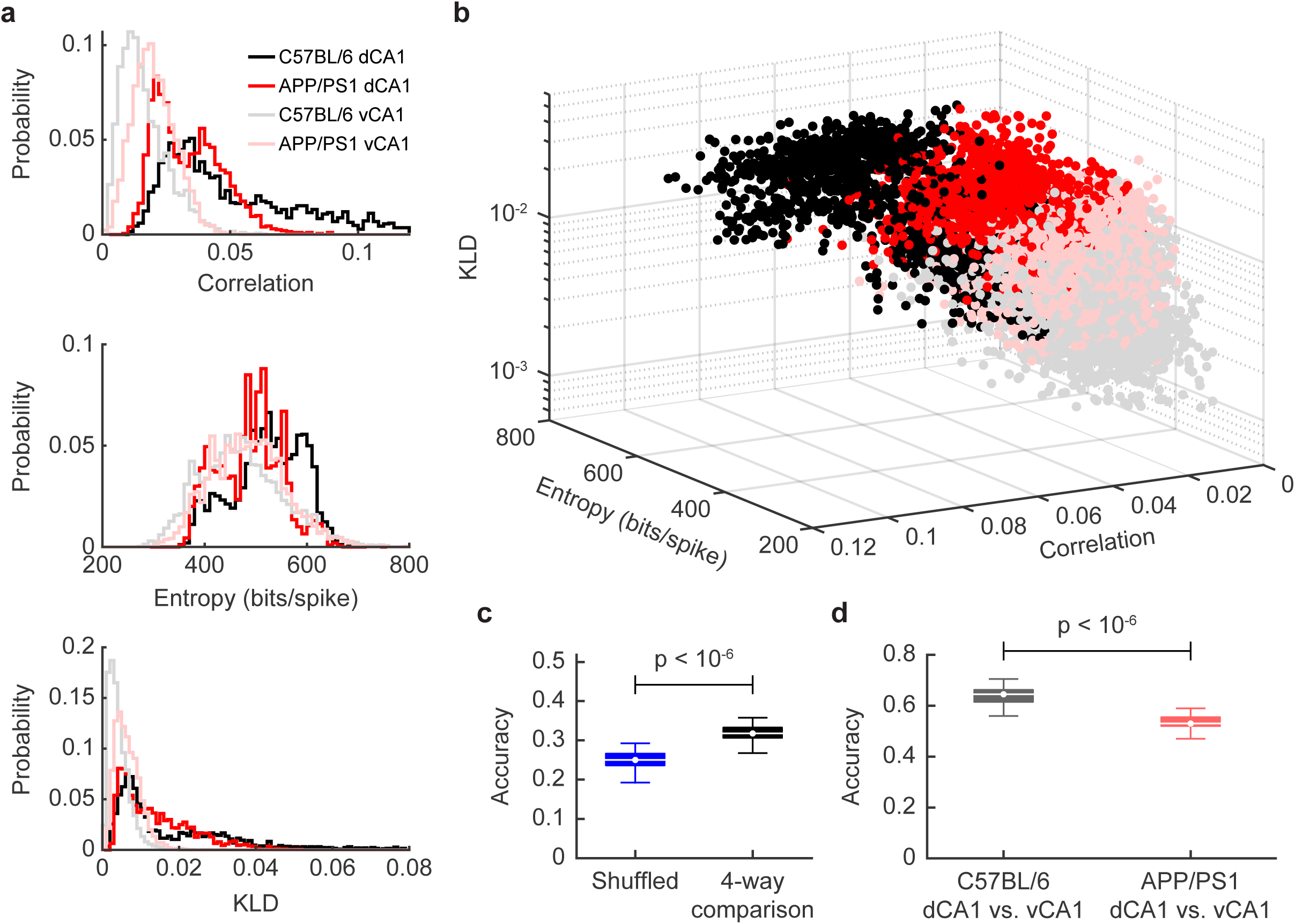
A signature of the differential impact of Aβ pathology on population activity is visible in the three-dimensional space defined by pairwise correlations, entropy, and the KLD of the maximum entropy model. (a) Distribution of correlation coefficients (top), entropy (middle), and KLD of the maximum entropy model (bottom) in dorsal and ventral CA1 in C57BL/6 and APP/PS1 mice (n_C57BL/6 dCA1_ = 2500 samples from 5 recording sessions, n_APP/PS1 dCA1_ = 2000 samples from 4 recording sessions, n_C57BL/6 vCA1_ = 3000 samples from 6 recording sessions, n_APP/PS1 vCA1_ = 4000 samples from 8 recording sessions). (b) Plot of correlations, entropy, and KLD of maximum entropy model for all recording sessions. Each point denotes a single resample (n_C57BL/6 dCA1_ = 2500 samples from 5 recording sessions, n_APP/PS1 dCA1_ = 2000 samples from 4 recording sessions, n_C57BL/6 vCA1_ = 3000 samples from 6 recording sessions, n_APP/PS1 vCA1_ = 4000 samples from 8 recording sessions). (c) Performance of decoder in a 4-way classification task to distinguish C57BL/6 dCA1, APP/PS1 dCA1, C57BL/6 vCA1, and APP/PS1 vCA1 samples (n = 100 iterations). (f) Performance of decoder in a 2-way classification task to distinguish C57BL/6 dCA1 from C57BL/6 vCA1 samples (n = 100 iterations). (d) Performance of decoder in a 2-way classification task to distinguish APP/PS1 dCA1 from APP/PS1 vCA1 samples (n = 100 iterations) p < 10^-4^, two-sided Wilcoxon rank-sum test.

To quantify these observations, we built a decoder to predict the strain identity and brain region of a subsample based on its population activity parameters (see Materials and Methods). The decoder performed significantly better than chance in a 4-way classification task (C57BL/6 dCA1 vs. APP/PS1 dCA1 vs. C57BL/6 vCA1 vs. APP/PS1 vCA1) (Figure 8c), suggesting that population activity can be used to distinguish both brain regions and mouse strains. Interestingly, however, the classifier was better able to distinguish dorsal and ventral CA1 in C57BL/6 mice than in APP/PS1 mice (Figure 8d). This suggests that in control mice, population activity varied across the longitudinal hippocampal axis, but that in APP/PS1 mice, this difference was degraded. In other words, Aβ pathology appears to have a homogenizing effect on population coding throughout the hippocampus, rendering the structure of network activity more similar in dorsal and ventral CA1. Taken together, these findings show that Aβ pathology distorts the geometry of the population code across the dorsoventral axis of the CA1 region of the hippocampus, and that the direction of that distortion varies depending on the hippocampal region.

## Discussion

We recorded the activity of large neuronal populations in awake C57Bl/6 and APP/PS1 mice across dorsal and ventral CA1 as the animals navigated virtual reality environment. Consistent with previous studies that examined spontaneous population activity in dCA1 in APP/PS1 animals (Chockanathan et al., 2020), key markers of population activity, such as correlations, entropy, and prediction error of a pairwise maximum entropy model, were reduced in APP/PS1 mice between 14-19 months of age while they were in a virtual reality environment. These suggest that the changes observed are a general feature of the Aβ pathology on circuits in dCA1. In vCA1, however, we found increased correlations, increased entropy, and a greater error in the maximum entropy model in APP/PS1 mice as compared to age matched control animals. These results show a differential impact of Aβ pathology on dorsal versus ventral CA1. Taken together, they suggest that the impact of AD on population activity is not simply a function of Aβ burden, but also a product of the interplay between histopathology and the properties of the neurons and circuits in each region.

The divergent impacts on population coding observed in the APP/PS1 mice across these two subregions could arise from multiple sources. First Aβ pathology could have a different effect on the neurons and intrinsic circuits of dCA1 as compared to those of vCA1 because these circuits are different. Differences across the longitudinal axis of the CA1 region include variations in gene expression (Cembrowski et al., 2016; Thompson et al., 2008), differences in neuronal morphology, and different biophysical properties in the CA1 pyramidal cells (Dougherty et al., 2012; Malik et al., 2016). For example, vCA1 neurons have lower dendritic length, decreased apical dendritic branching, higher membrane potential, and higher input resistance than dCA1 neurons (Dougherty et al., 2012; Malik et al., 2016). As a result of these differences, the structure of population activity would be different, something we found in this work. A specific pathology then, such as Aβ accumulation, would then exert divergent effects on different parts of the hippocampus because the networks in these regions are functionally different.

A second possibility is that Aβ pathology differentially impacts the brain regions that project to either dorsal or ventral CA1. There exist substantial variations in afferent innervation throughout the longitudinal hippocampal axis. For example, there is a topographic organization of entorhinal cortex (EC)-hippocampus circuits, such that dorsal hippocampus preferentially receives projections from dorsolateral EC, while ventral hippocampus preferentially receives input from ventromedial EC (Dolorfo and Amaral, 1998; van Strien et al., 2009; Witter et al., 2000). In turn, each of these EC subregions have their own distinct patterns of afferent connectivity; for example, dorsal and lateral EC receive inputs from anterior cingulate and retrosplenial cortex while ventral and medial EC receive inputs from infralimbic and prelimbic cortex (Jones and Witter, 2007; Strange et al., 2014; Witter, 1993). In addition, a recent study of direct whole-brain inputs to CA1 projection neurons (PNs) showed that vCA1 PNs receive a more diverse array of inputs from throughout the brain, including the basolateral amygdala and piriform cortex, than dCA1 PNs, whose inputs primarily arise from the EC and other hippocampal subfields (Tao et al., 2021). Given the variations in Aβ burden across different brain regions (Liebmann et al., 2016; Whitesell et al., 2018), it is possible that the differences in population activity between dorsal and ventral CA1 are inherited from upstream circuits.

Our work suggests that the differences in network activity that exist across the longitudinal axis of the hippocampus (the initial conditions of the circuit) are differentially altered by the Aβ accumulation in the APP/PS1 animals. A natural question is how might a single pathology exert such heterogeneous, often opposite effects on the networks of the hippocampus? Evidence suggests that the effects of Aβ on neurons and synapses can be heterogeneous, with the precise nature of the disruption complex and contingent on many factors (Busche et al., 2008; Palop and Mucke, 2010). For example, moderate elevations in Aβ appear to facilitate synaptic potentiation (including long-term potentiation (LTP)) by increasing presynaptic release probabilities (Abramov et al., 2009b; Puzzo et al., 2008), while large increases in Aβ levels lead to decreased LTP and enhanced long term depression (LTD) by altering post-synaptic mGluR and NMDAR signaling (Li et al., 2009; Puzzo et al., 2008). Coupled with findings of differences in the induction and magnitude of LTP and LTD between dorsal and ventral CA1 neurons (Kouvaros and Papatheodoropoulos, 2016; Maggio and Segal, 2009; Maruki et al., 2001; Milior et al., 2016), these observations hint at the possibility that Aβ pathology may differentially affect synaptic plasticity in dorsal and ventral hippocampus. Multiple studies have shown that Aβ pathology was associated with deficits in the induction or maintenance of LTP in the dorsal hippocampus (Chapman et al., 1999; Larson et al., 1999; Moechars et al., 1999). Changes in synaptic coupling due to the differential effects of Aβ pathology in the APP/PS1 mice could induce either LTP or LTD across different cell types and different regions of the hippocampus, leading to diverse impacts on the correlations that would arise from changes in synaptic strength. Furthermore, studies showing differences in the intrinsic excitability of neurons in mouse models of Aβ pathology (Šišková et al., 2014) also suggest that the biophysical properties of neurons that influence spike-timing dependent plasticity (Bi and Poo, 1998; Froemke et al., 2005; Froemke and Dan, 2002) could either increase or decrease correlations depending on whether Aβ renders those neurons more excitable or less excitable. In this regard, the diametric effects of Aβ pathology in the APP/PS1 animals on correlations, entropy, and the structure of population activity may reflect the heterogeneous effects of that pathology on individual neurons and synapses.

We consider what this result might tell us about how network activity determines behavior. In patients with AD, many cognitive domains (episodic memory, spatial cognition, etc.) are blunted (Appell et al., 1982; Greene et al., 1996; Monacelli et al., 2003; Morris and Kopelman, 1986; Murdoch et al., 1987), while other domains, (stress, anxiety, etc.), have been shown to be enhanced (Apostolova and Cummings, 2008; Hwang et al., 2004; Mega et al., 1996; Peters et al., 2012). The findings of this study offer one insight into a potential mechanism behind these disparate effects. In an ensemble coding scheme, a reduction in entropy or in pairwise correlations would erode representations, something that might give rise to loss in a cognitive or mnemonic domain. By contrast, increases in correlations and entropy may provide a neural substrate for enhanced anxiety or fear. Correlations and increased entropy could be thought of as increases in generalization, which would, for example, alter how sensory experiences are mapped onto anxiety. Our results suggest that understanding the behavioral changes in AD may emerge not only by looking at cellular pathology, but the diverse impact that pathology has on network function including the ways in which neuronal population activity is altered.

## Supplemental figure legends

**Figure 1-figure supplement 1:**
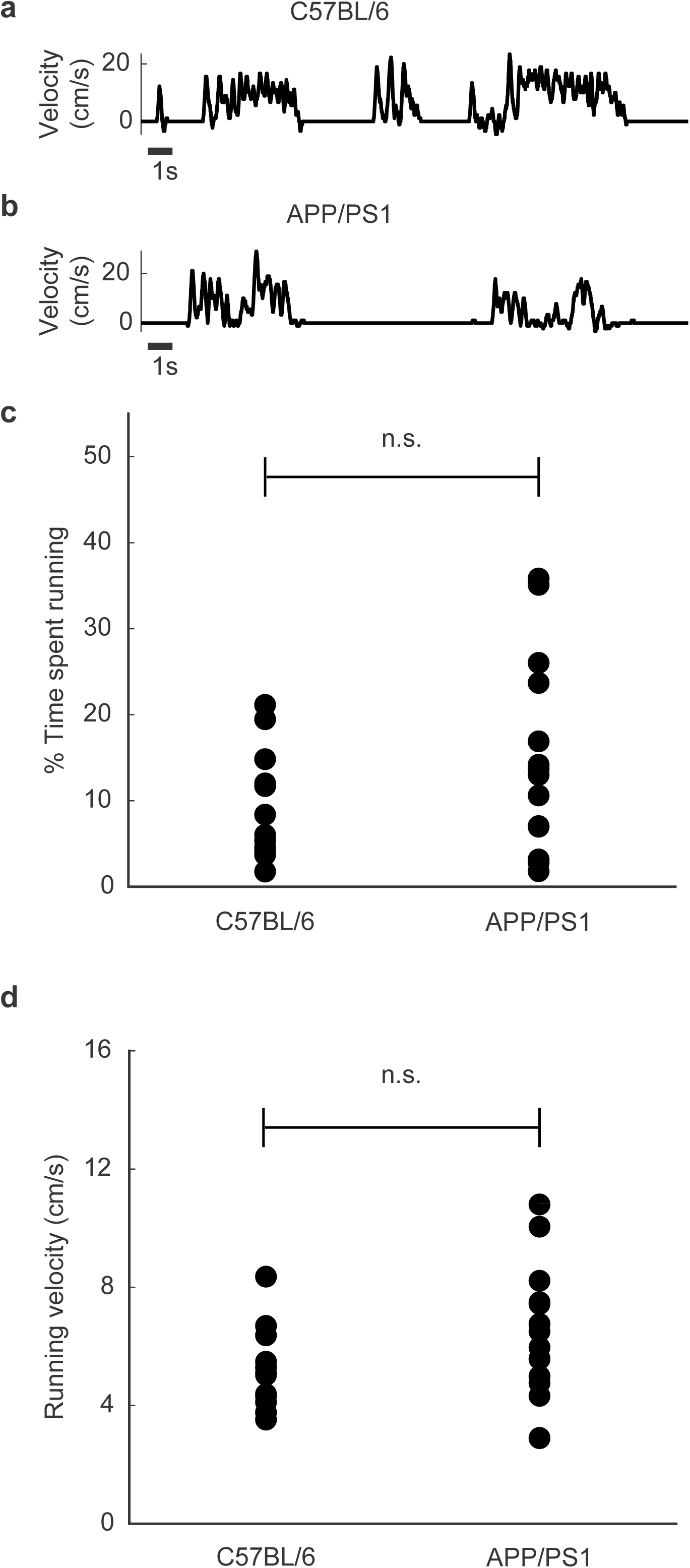
Running behavior was similar between C57BL/6 and APP/PS1 mice. (a) Representative trace of running velocity for a C57BL/6 mouse. (b) Representative trace of running velocity for an APP/PS1 mouse. (c) There was no significant difference in the proportion of time spent running between the C57BL/6 and APP/PS1 mice (mean ± std: C57BL/6 = 9.4 ± 6.4%, APP/PS1 = 14.9 ± 10.9%, p = 0.21, two-sided Wilcoxon sign-rank test, n = 6 animals). Each point denotes a single animal. (d) There was no significant difference in the mean running velocity of the C57BL/6 and APP/PS1 mice (mean ± std: C57BL/6 = 5.2 ± 1.4 cm/s, APP/PS1 = 6.3 ± 2.3 cm/s, p = 0.16, two-sided Wilcoxon sign-rank test, n = 6 animals). Each point denotes a recording session.

**Figure 4-figure supplement 1:**
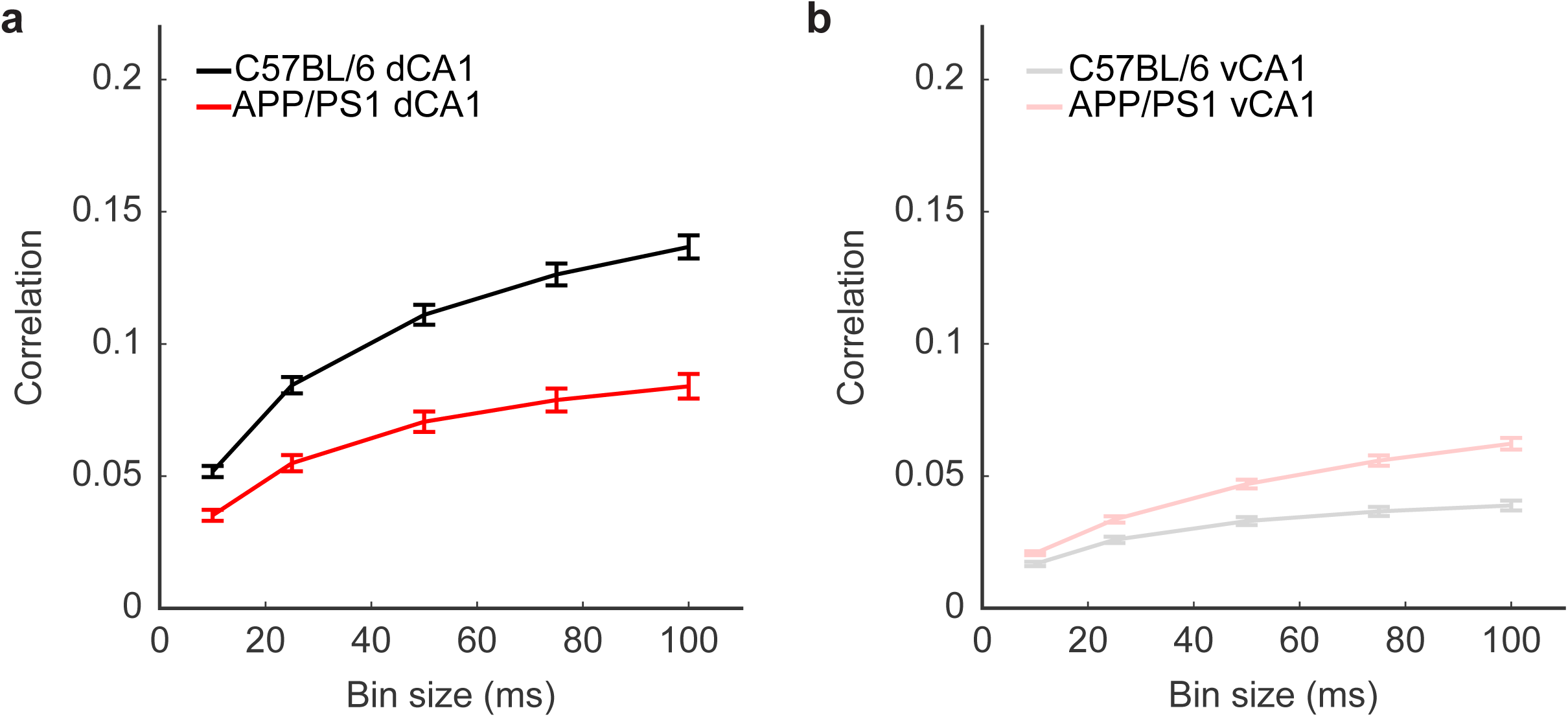
Differences in pairwise correlation were preserved across a range of temporal bin sizes. (a) In dCA1, APP/PS1 mice had significantly lower correlations than C57BL/6 mice when spiking activity was binned at 5ms, 25ms, 50ms, 75ms, and 100ms (for all bin sizes, p < 10^-6^, two-sided Wilcoxon rank-sum test, n_C57BL/6_ = 1250 unit pairs from 5 recording sessions, n_APP/PS1_ = 1000 unit pairs from 4 recording sessions). Error bars show the standard error of the mean. (b) In vCA1, APP/PS1 mice had significantly higher correlations than C57BL/6 mice when spiking activity was binned at 5ms, 25ms, 50ms, 75ms, and 100ms (for all bin sizes, p < 10^-6^, two-sided Wilcoxon rank-sum test, n_C57BL/6_ = 1500 unit pairs from 6 recording sessions, n_APP/PS1_ = 2000 unit pairs from 8 recording sessions). Error bars show the standard error of the mean.

**Figure 4-figure supplement 2:**
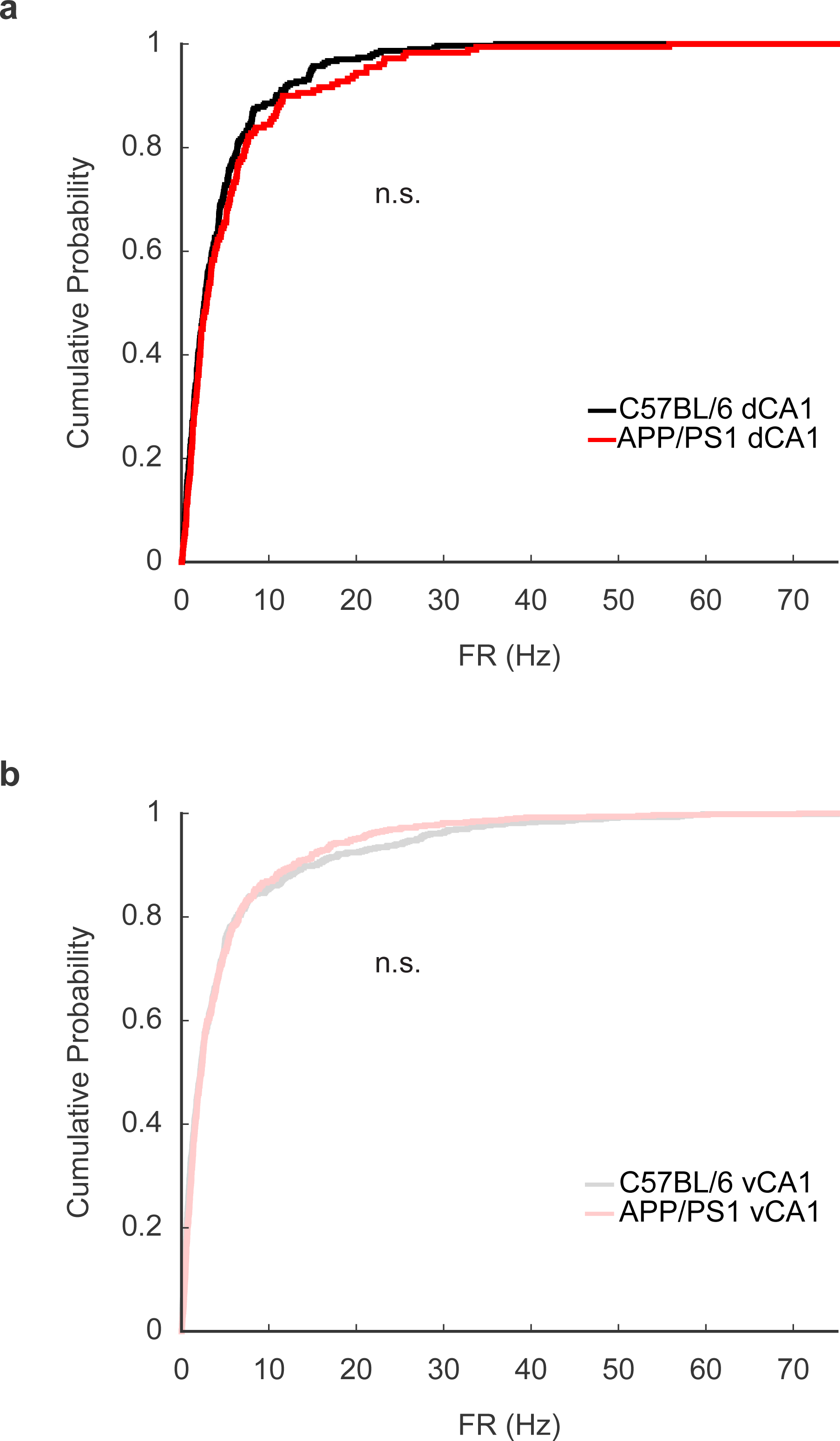
Similar mean firing rates between C57BL/6 and APP/PS1 mice. (a) In dCA1, firing rates in C57BL/6 and APP/PS1 mice were not significantly different (mean ± std: C57BL/6 = 4.4 ± 5.3 Hz, APP/PS1 = 5.4 ± 7.3 Hz, p = 0.43, two-sided Wilcoxon rank-sum test, n_C57BL/6_ = 305 neurons, n_APP/PS1_ = 180 neurons). (b) In vCA1, firing rates in C57BL/6 and APP/PS1 mice were not significantly different (mean ± std: C57BL/6 = 5.5 ± 9.5 Hz, APP/PS1 = 4.9 ± 7.8 Hz, p = 0.78, two-sided Wilcoxon rank-sum test, n_C57BL/6_ =689 neurons, n_APP/PS1_ = 651 neurons).

**Figure 5-figure supplement 1:**
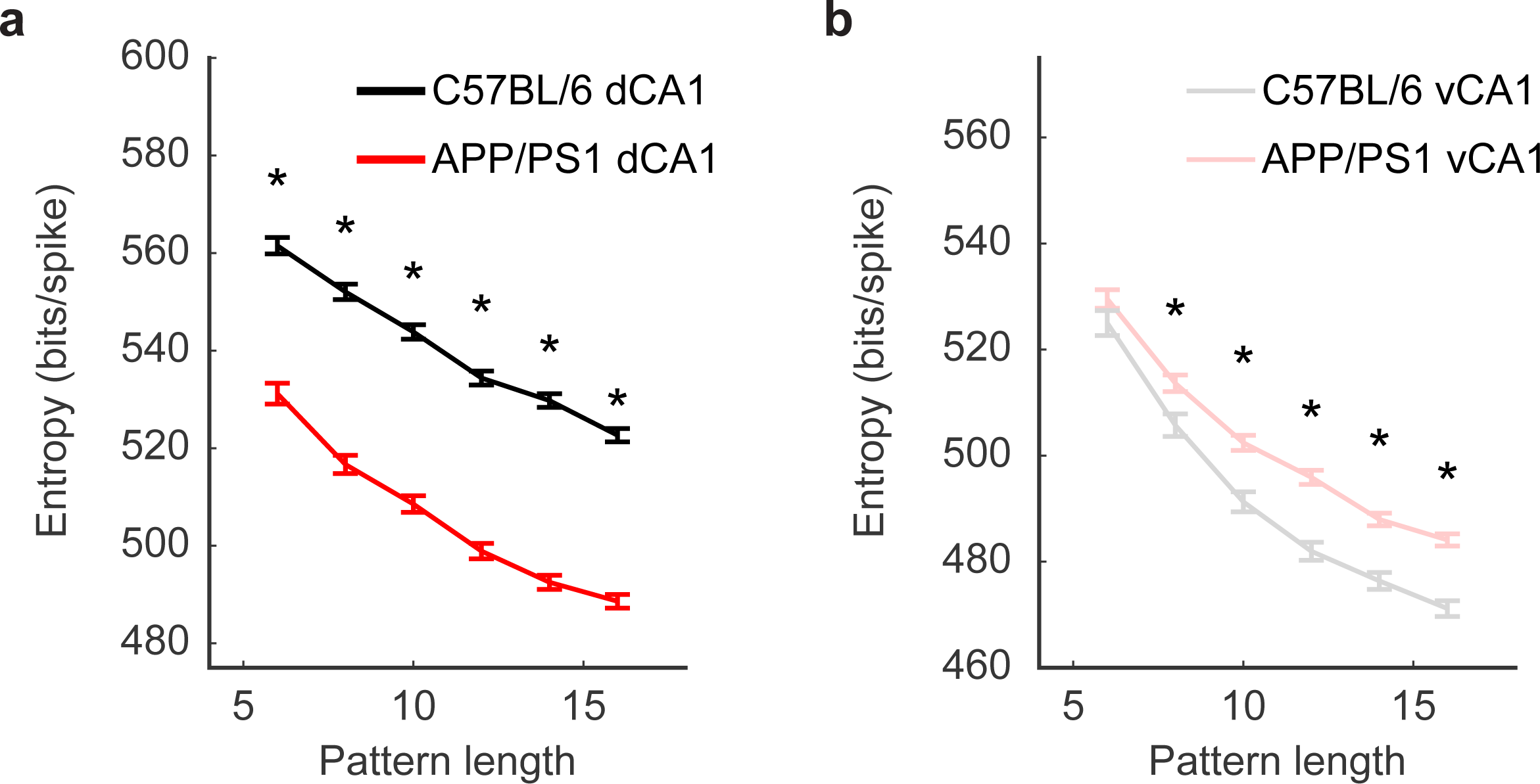
Differences in population entropy were preserved across a range of pattern subpopulation sizes. (a) In dCA1, APP/PS1 mice had a significantly lower entropy with subpopulation sizes of 6, 8, 10, 12, 14, and 16 (asterisks denote p < 10^-6^, two-sided Wilcoxon rank-sum test, n_C57BL/6_ = 2500 subsamples from 5 recording sessions, n_APP/PS1_ = 2000 subsamples from 4 recording sessions). Error bars show the standard error of the mean. (b) in vCA1, APP/PS1 mice had a significantly higher entropy with subpopulation sizes of 8, 10, 12, 14, and 16 (asterisks denote p < 10^-3^, two-sided Wilcoxon rank-sum test, n_C57BL/6_ = 3000 subsamples from 6 recording sessions, n_APP/PS1_ = 4000 subsamples from 8 recording sessions). Error bars show the standard error of the mean.

**Figure 6-figure supplement 1:**
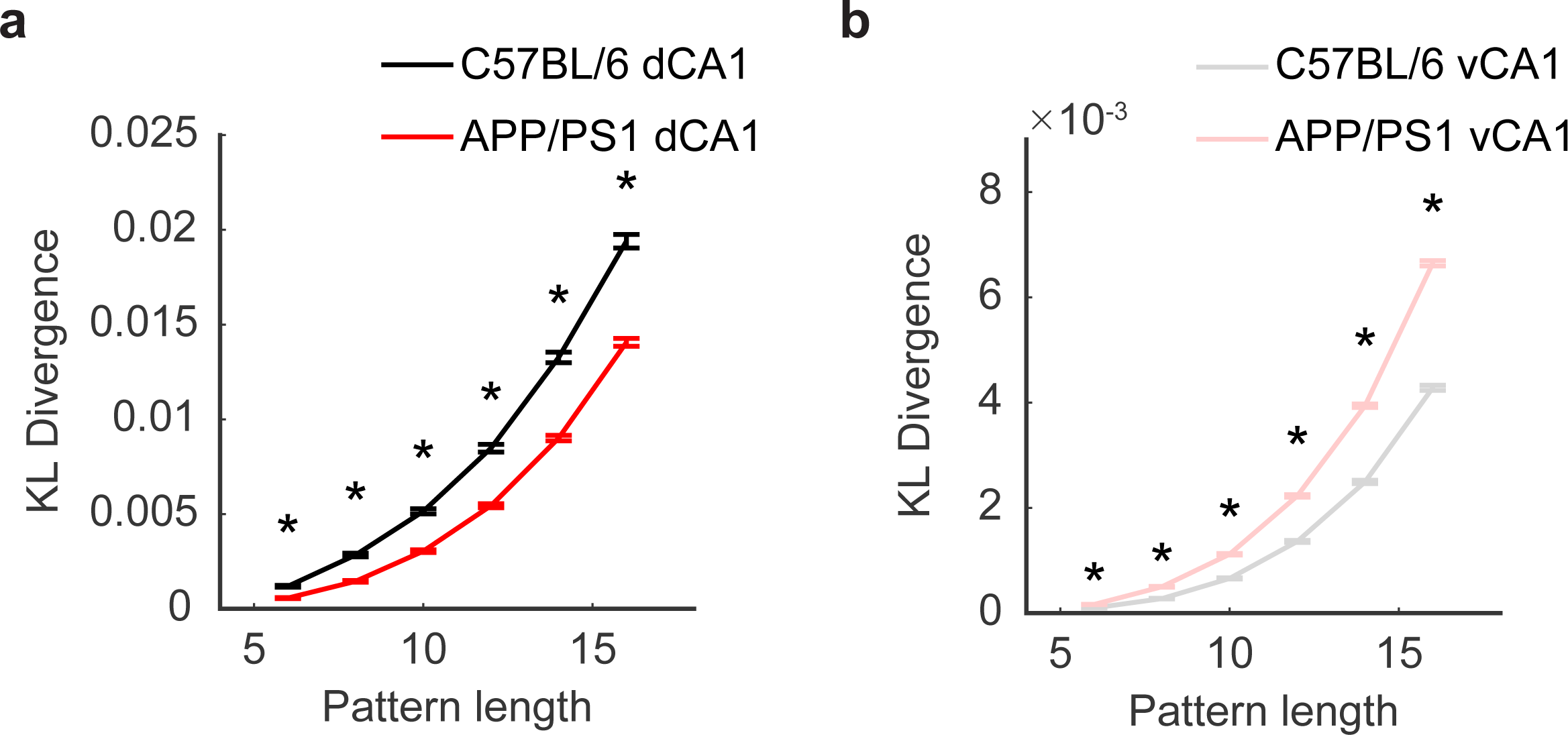
Differences in maximum entropy model KLD were preserved across a range of pattern subpopulation sizes. (a) In dCA1, APP/PS1 mice had a significantly lower entropy with subpopulation sizes of 6, 8, 10, 12, 14, and 16 (asterisks denote p < 10^-4^, two-sided Wilcoxon rank-sum test, n_C57BL/6_ = 2500 subsamples from 5 recording sessions, n_APP/PS1_ = 2000 subsamples from 4 recording sessions). Error bars show the standard error of the mean. (b) In vCA1, APP/PS1 mice had a significantly higher entropy with subpopulation sizes of 6, 8, 10, 12, 14, and 16 (asterisks denote p < 10^-6^, two-sided Wilcoxon rank-sum test, n_C57BL/6_ = 3000 subsamples from 6 recording sessions, n_APP/PS1_ = 4000 subsamples from 8 recording sessions). Error bars show the standard error of the mean.

**Figure 7-figure supplement 1:**
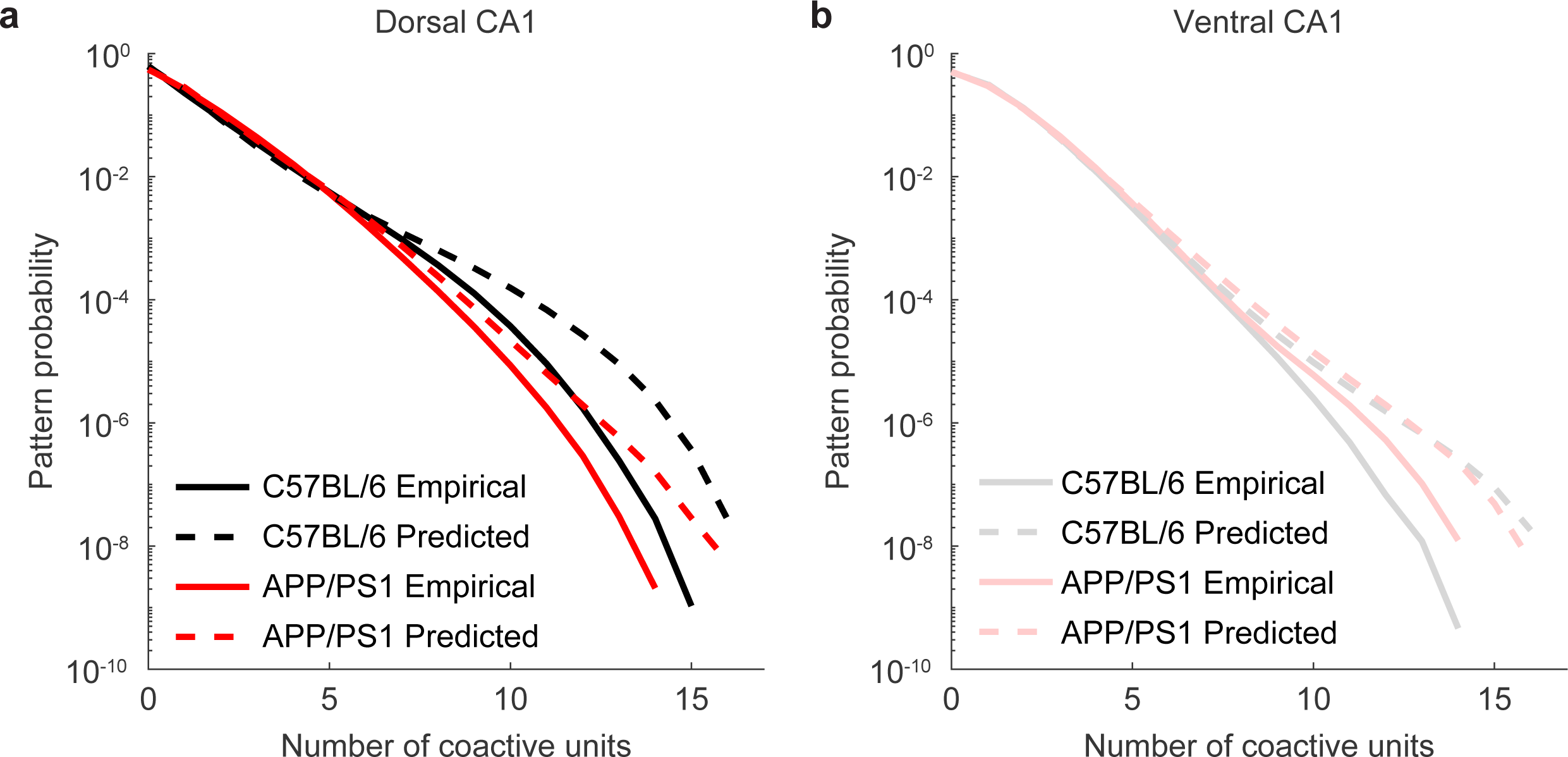
Maximum entropy model predicted probabilities and empirical probabilities for patterns grouped by number of coactive units. In both dorsal and ventral CA1, the difference between the empirical and predicted probabilities is larger for patterns with larger numbers of coactive units. (a) In dCA1, the difference predicted and empirical pattern probabilities for APP/PS1 mice is smaller, implying a better prediction, than for C57BL/6 mice. Pattern probabilities were averaged across 2500 subsamples from 5 recording sessions in C57BL/6 mice and across 2000 subsamples from 4 recording sessions in APP/PS1 mice. (b) In vCA1, the difference between the predicted and empirical pattern probabilities for APP/PS1 mice is larger, implying a worse prediction, than for C57BL/6 mice. Pattern probabilities were averaged across 3000 samples from 6 recording sessions in C57BL/6 mice and across 4000 samples from 8 recording sessions in APP/PS1 mice.

## Acknowledgements

We thank Daniel Guarino and Martin Gira at the University of Rochester Center for Visual Science for their assistance with constructing the virtual reality platform used in this experiment. We also thank Jeff Fox at the University of Rochester Center for Musculoskeletal research for his technical assistance with the slide scanner and image acquisitions. K.P. was supported by the National Science Foundation CAREER Grant 1749772, the National Institute of Mental Health Grant R01MH11392, the Schmitt Foundation, and the Cystinosis Research Foundation. U.C. was supported by the National Institute of General Medical Sciences Grant T32 GM007356.

